# Grid cell firing fields in a volumetric space

**DOI:** 10.1101/2020.12.06.413542

**Authors:** Roddy M. Grieves, Selim Jedidi-Ayoub, Karyna Mishchanchuk, Anyi Liu, Sophie Renaudineau, Éléonore Duvelle, Kate J. Jeffery

## Abstract

We investigated how entorhinal grid cells represent volumetric (three-dimensional) space. On a flat surface, grid cell firing fields are circular and arranged in a close-packed hexagonal array. In three dimensions, theoretical and computational work suggests that the most efficient configuration would be a regular close packing of spherical fields. We report that in rats exploring a cubic lattice, grid cells were spatially stable and maintained normal directional modulation, theta modulation and spike dynamics. However, while the majority of grid fields were spherical, they were irregularly arranged, even when only fields abutting the lower surface (equivalent to the floor) were considered. Thus, grid organization is shaped by the environment’s movement affordances, and may not need to be regular to support spatial computations.

**One Sentence Summary:** In rats exploring a volumetric space, grid cells are spatially modulated but their firing fields are irregularly arranged.

## Main Text

Entorhinal grid cells tile an environment’s surface with evenly spaced firing fields that provide a metric supporting the brain’s spatial cognitive map (*1*). An unresolved but central question is whether this map is three-dimensional, as befits the behavioral ecology of most vertebrates. Hippocampal place cells, the core of the cognitive map in mammals (*2*) and probably birds (*3*) form spatially defined firing fields in a volumetric lattice in both bats (*4, 5*) and rats (*6*), suggesting a capacity for the vertebrate brain to fully map volumetric space. We investigated whether this map is founded on a three-dimensional entorhinal grid.

Theoretical considerations suggest that in a volumetric space, an optimal grid structure for 3D spatial mapping would be a hexagonal-close-packed (HCP) or face-centered-cubic (FCC) lattice of firing fields (Fig. 1A, *7, 8*). However, previous studies on vertical surfaces found that grid fields formed stripes (*9*) or expanded blobs (*10*) depending on the locomotor affordances (potential for movement) of the surface. Here, using wireless telemetry on rats exploring a cubic lattice maze, we investigated whether grid fields are indeed close-packed (i.e., optimally organized). We show that grid cells stably express spatial, spheroid fields but in a random pattern, and explore the implications of this for spatial computations.

**Fig. 1:**
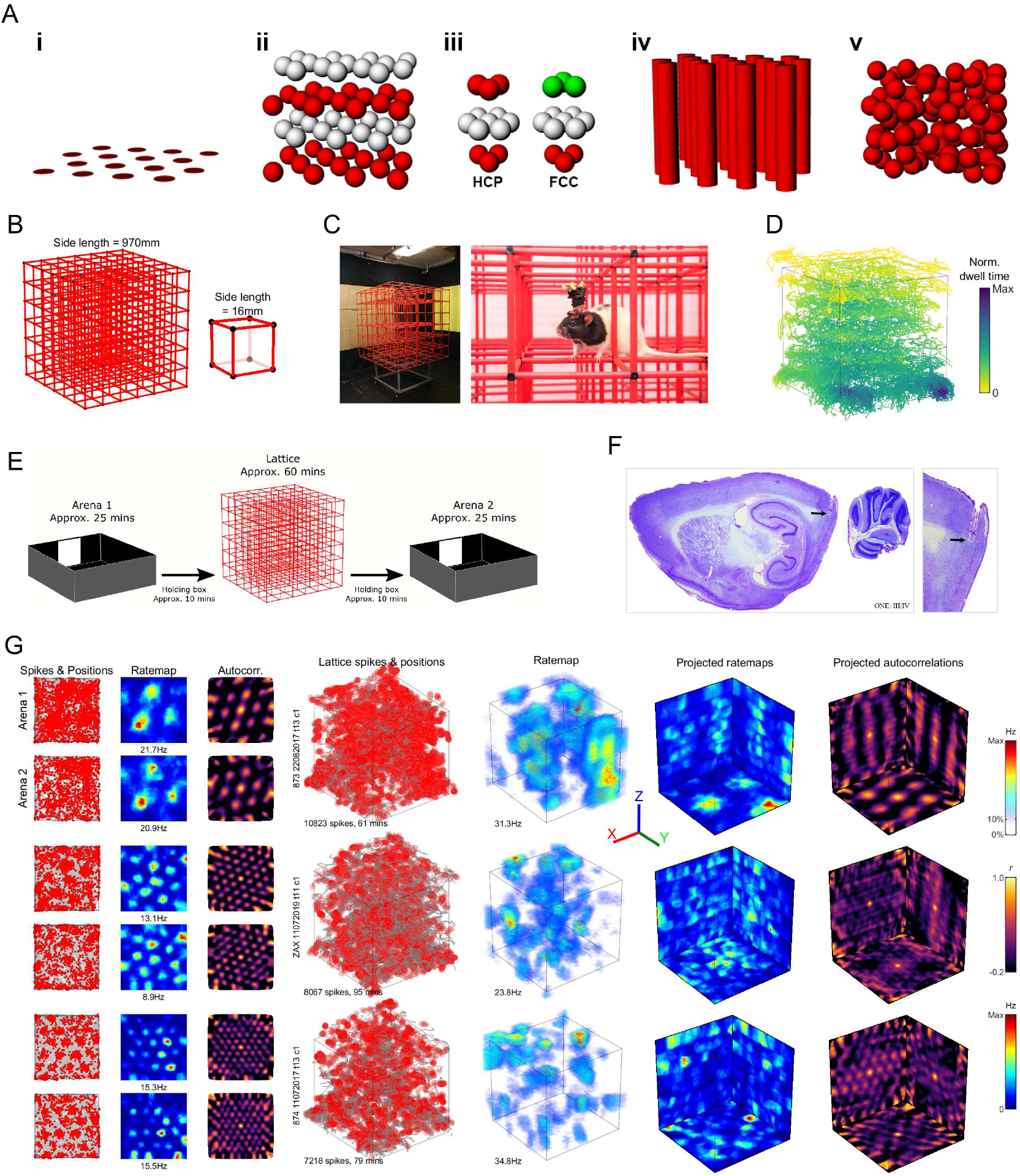
Grid cells produced firing fields in a three-dimensional climbing lattice. (A) Hypothetical grid field packings. i) Standard horizontal hexagonal field configuration; ii) Exploded close-packed lattice, in this case HCP (layers color-coded for clarity); iii) Units of the two optimal packings: HCP (left) alternates two layer-arrangements while FCC (right) has three; iv) columnar field configuration; v) random field configuration. (B) Lattice maze schematic. (C) Lattice maze photographs. (D) Example coverage in a lattice session. Color denotes normalized dwell time in each region. (E) Recording protocol. (F) Example histology. (G) Three representative grid cells in the arenas (left) and lattice (right). Left-right: arena spike plots (gray = coverage; red dots = spikes), arena ratemaps, arena autocorrelations, volumetric spike plots, volumetric firing rate maps, rate maps as projected onto each of the three coordinate planes, and projected autocorrelations. Colorbars from top to bottom correspond to volumetric ratemaps, autocorrelations and planar ratemaps.

We recorded medial entorhinal cortex (mEC) grid cells in 7 rats freely foraging within a 3D lattice maze (*6, 11–13*)(approximately 1m^3^; Fig. 1B to C) and a standard horizontal arena (1.2m × 1.2m; Fig. 1E & Fig. S1A). Rats fully explored the environments but spent more time in the bottom layer of the lattice (Fig. 1D; Fig. S1B-E). They mainly moved parallel to the maze boundaries and prioritized horizontal movements (Fig. S1F; see (*13*) for in-depth behavior analysis).

We recorded a total of 47 grid cells in layers I-IV of the mEC (Fig. 1F to G, Fig. S2 for all grid cells, Movie S1 for rotating plots, Fig. S3 for all histology, Table S1 for per rat summary). Grid cells maintained the same firing rates in the arena and lattice (Fig. S4A). Grid cells were stable throughout recording as shown by high grid scores in each arena (a measure of hexagonality) and the cross-correlation between them (Fig. S4B-D) in combination with high correlations between arena maps (Fig. S4E).

In both mazes grid cell firing was stable between session halves (session halves vs shuffled: *p* < .001 in all cases, one-sample t-tests; no difference between mazes: *p* = .20, one-way ANOVA; Fig. 2A); with increased stability in the XY plane of the lattice (Fig. S5B). Spatial information was also higher than spike-train-shuffled data (*p* < .05 in all cases, one-sample t-tests), indicating that firing was more spatially clustered than chance (Fig. 2B) but this was closer to chance in the lattice (*F*(2,137) = 20.3, *p* < .0001, *η*^2^ = 0.228, lattice vs arena 1 or 2: *p* < .0001, all other: *p* > .05, one-way ANOVA; Fig. 2B) and there was a positive correlation between arena and lattice spatial information (Fig. 2C).

**Fig. 2:**
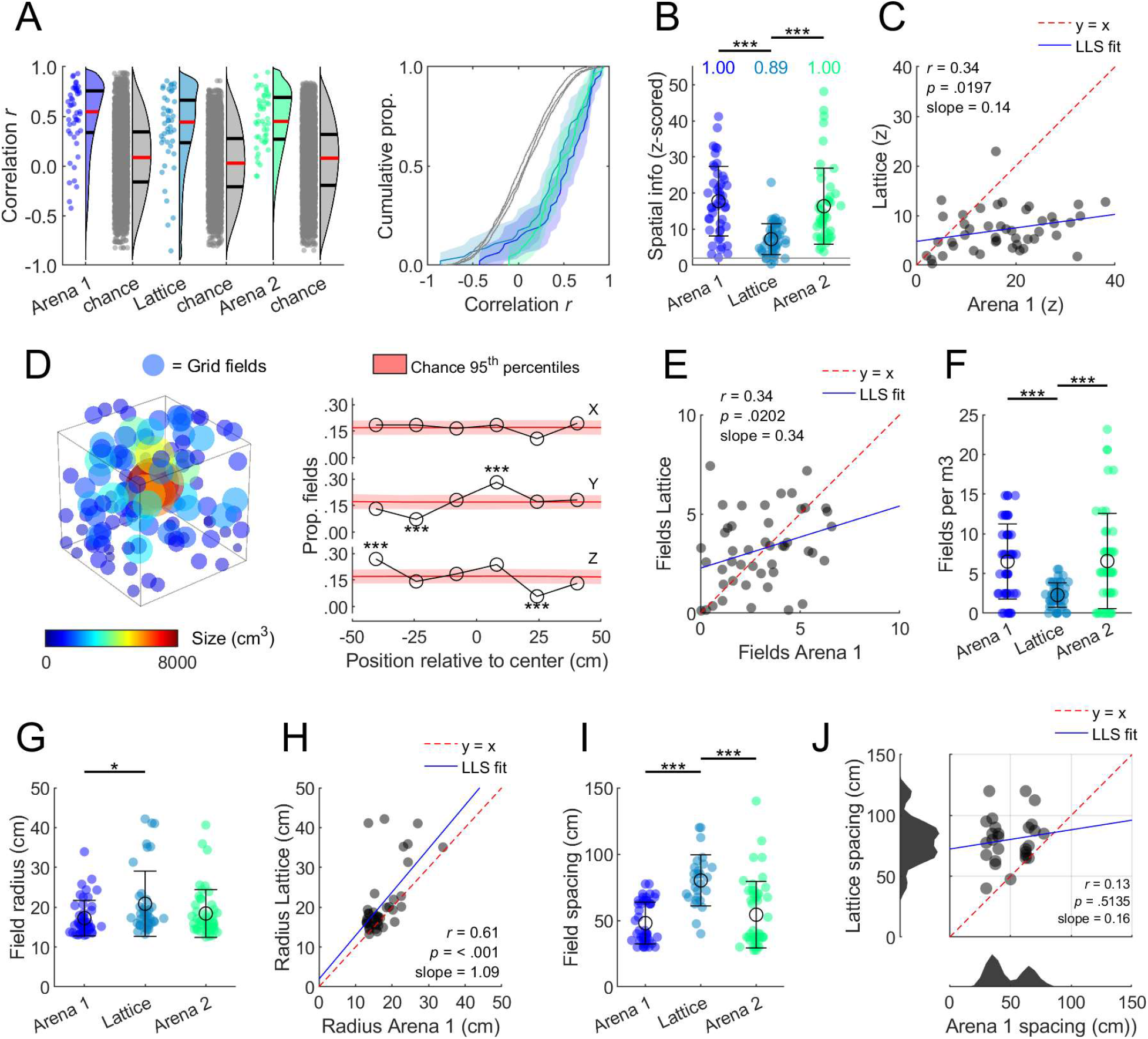
Grid cells mapped the lattice with large and widely spaced but stable fields. (A) Grid cell firing was spatially correlated between session halves (red lines denote medians; black lines denote 1^st^ and 3^rd^ quantiles. (B) Z-scored spatial information was higher than chance in all environments but reduced in the lattice; text gives the proportion of cells exceeding the shuffle 95^th^ percentile (z = 1.96; gray line). (C) Arena and lattice spatial information was significantly positively correlated. LLS: linear least squares line fit. (D) Position and size of every grid field (left) and proportion of fields in every lattice layer (right). (E) The number of fields per grid cell in the arena and lattice was positively correlated. (F) Grid cells exhibited significantly fewer fields per m^3^ in the lattice maze. (G) Grid field radius was significantly larger in the lattice than the first arena. (H) The size of grid fields in the arena and lattice was significantly positively correlated. (I) Grid spacing was significantly larger in the lattice. (J) Grid spacing (max. 120 cm) in the arena and lattice was uncorrelated, and arena grid modules (bottom histogram) were disrupted in the lattice. Cells for which no lattice spacing could be estimated (40.2%) are not shown in I or J. See Fig. S6 for schematic and validation of procedures in G-J.

Grid fields were distributed throughout the lattice volume (Fig. 2D). The number of fields expressed in the arena and lattice was significantly positively correlated (Fig. 2E). Surprisingly, the number of fields exhibited in the arena and lattice did not differ (arena 1: 2.6 ±0.28; lattice: 2.9 ±0.29; arena 2: 2.6 ±0.34 mean ±SEM fields; *F*(2,137) = 0.3, *p* = .72), resulting in significantly fewer grid fields per m^3^ in the lattice (*F*(2,137) = 13.8, *p* < .0001, *η*^2^ = 0.167, lattice vs arena 1 or 2 *p* < .0001, all other *p* > .05, one-way ANOVA; Fig. 2F) which is consistent with hippocampal place cells (*6, 10*). Grid field radius was significantly larger in the lattice compared to the first arena (*F*(2,131) = 3.7, *p* = .0284, *η*^2^ = 0.053, arena 1 vs lattice *p* = .0254, all other *p* > .05, one-way ANOVA; Fig. 2G) and there was a significant positive correlation between the two (Fig. 2H). Grid cells also exhibited significantly larger spacing in the lattice (*F*(2,118) = 22.0, p < .0001, *η*^2^ = 0.271, lattice vs arena 1 or 2 *p* < .0001, all other *p* > .05, one-way ANOVA; Fig. 2I) but this was not correlated with arena spacing; instead, arena grid scale modules were disrupted in the lattice (Fig. 2J).

We next looked at the spatial pattern of firing fields in the lattice maze. Previous theoretical (*7, 8, 14*) and computational (*15*) work suggests that the optimal packing of grid fields in three-dimensions is a hexagonal close-packed (HCP) or face centered-cubic (FCC) configuration (Fig. 1A, *16, 17*). To test this, we first calculated a close-packed quality score (χ_CP_) that measures the presence of either an FCC or HCP structure. This was significantly lower than expected for an FCC or HCP arrangement and was instead comparable to columnar or random fields (Fig. 3A; *F*(4,442) = 1612.2, *p* < .0001, *η*^2^ = 0.936; all groups differ *p* < .001 except grid cells and random *p* > .05). Configuration-specific scores (for FCC, HCP and columns (COL); Fig. S7 and Fig. S8) were all close to zero and significantly lower than simulated configurations (Fig. 3B&C; Fig. S9). In simulated data the field configurations most similar to real data were the uniformly random (Fig. 3D) or shuffled ones (Fig. S10).

**Fig. 3:**
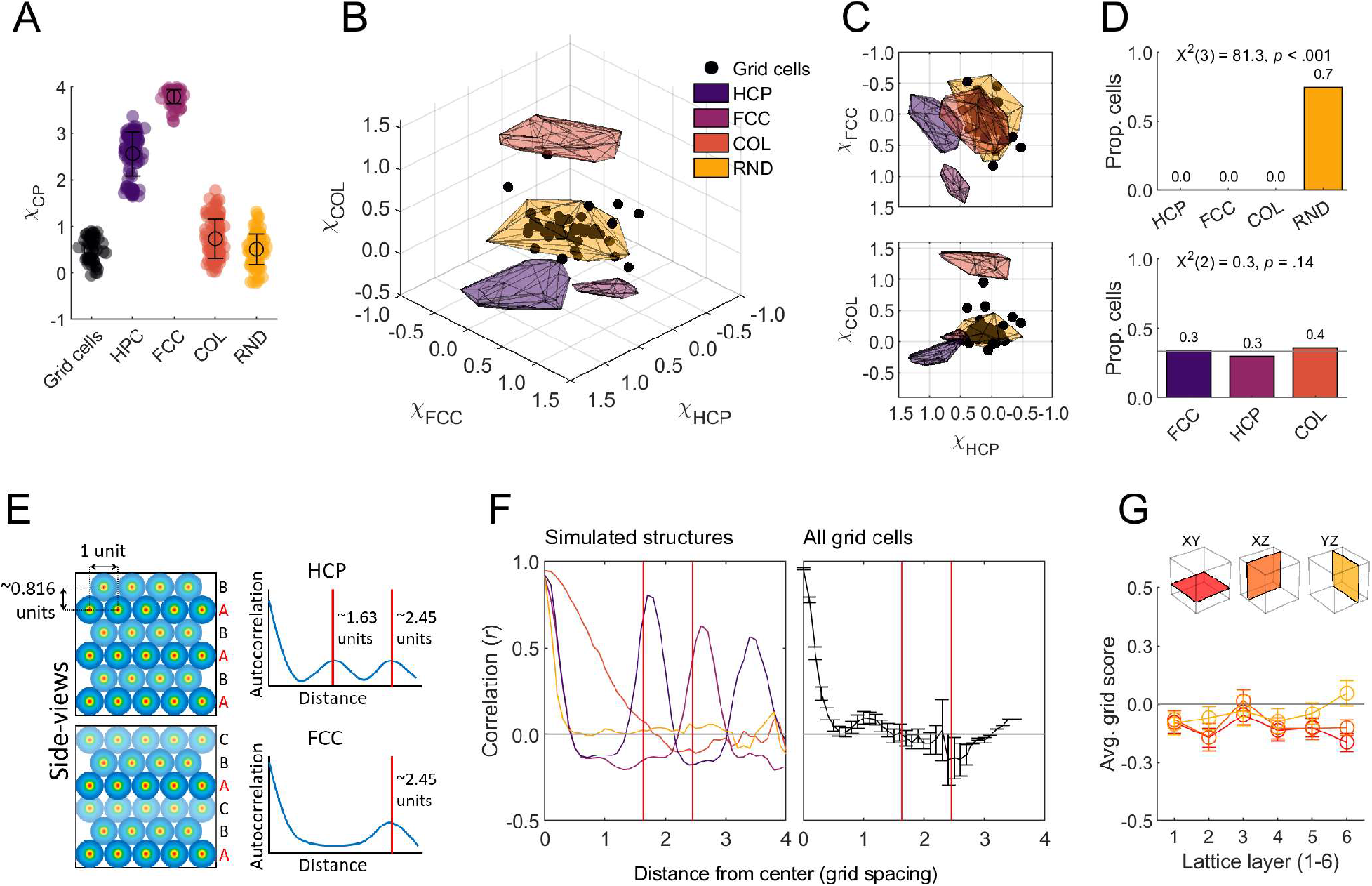
Grid fields were randomly distributed in the lattice. (A) Quality scores (χ_CP_) for grid cells vs. simulated configurations. (B) Structure scores (χ_FCC_, χ_HCP_ and χ_COL_) for grid cells (markers) and simulations (convex hulls shown as shaded polygons). (C) Two-dimensional side projections of B showing that grid cells overlapped the most with random configuration scores. (D) Grid cells categorized based on which convex hull they fell into (top; 30% of cells are uncategorized) or which configuration score was maximal (bottom). (E) Schematic of simulated HCP (top) and FCC (bottom) arrangements. Distance between repetitions of the same layer differ between layer repetitions in HCP (top) and FCC (bottom). (F) This analysis finds the expected peak correlation patterns in simulated configurations (left; colors same as A) but not real grid cells (right). (G) Grid cells exhibited low grid scores in all Cartesian coordinate planes of the lattice.

If a cell expressed an FCC or HCP firing pattern in the lattice its firing rate map should be periodically self-similar (i.e., correlate highly with itself) with vertical shifts at multiples of approximately 0.816*d*, where *d* is the cell’s grid spacing. Furthermore, the position of these similarity peaks would depend on the firing pattern (Fig. 3E). While this was detected in our simulations, grid cells did not show evidence of either pattern (Fig. 3F). Additionally, grid scores were generally negative for all layers of the lattice (Fig. 3G) and equal to chance for non-aligned planes (fig. S9E).

Grid fields were significantly more elongated in the lattice than the arena (Fig. 4A; *F*(2,374) = 48.2, *p* < .0001, *η*^2^ = 0.205, lattice vs arena 1 or 2 *p* < .0001, all other *p* > .05, one-way ANOVA) mainly along the vertical axis (Fig. 4B). After correcting for this field anisotropy grid cells still exhibited a random field configuration (Fig. S10A&B). Grid scores were significantly lower in all lattice projections when compared to the arena XY plane (Fig. 4C; *F*(3,181) = 169.6, *p* < .0001, *η*^2^ = 0.738, all lattice projections vs arena XY *p* < .0001, lattice XY vs XZ *p* = .0425, all other *p* > .05). However, a number of cells exhibited higher grid scores than expected by chance in the XY plane (12.8%; Fig. 4C&D). These cells were recorded in the same rat across multiple sessions and tetrodes. Evidence for square firing patterns (*17*) was also observed in some grid cells but the overall proportion was close to that expected by chance (6.5% for XY plane; Fig. S12A&B) and scores were similar to those of non-grid cells (Fig. S12C&D).

**Fig. 4:**
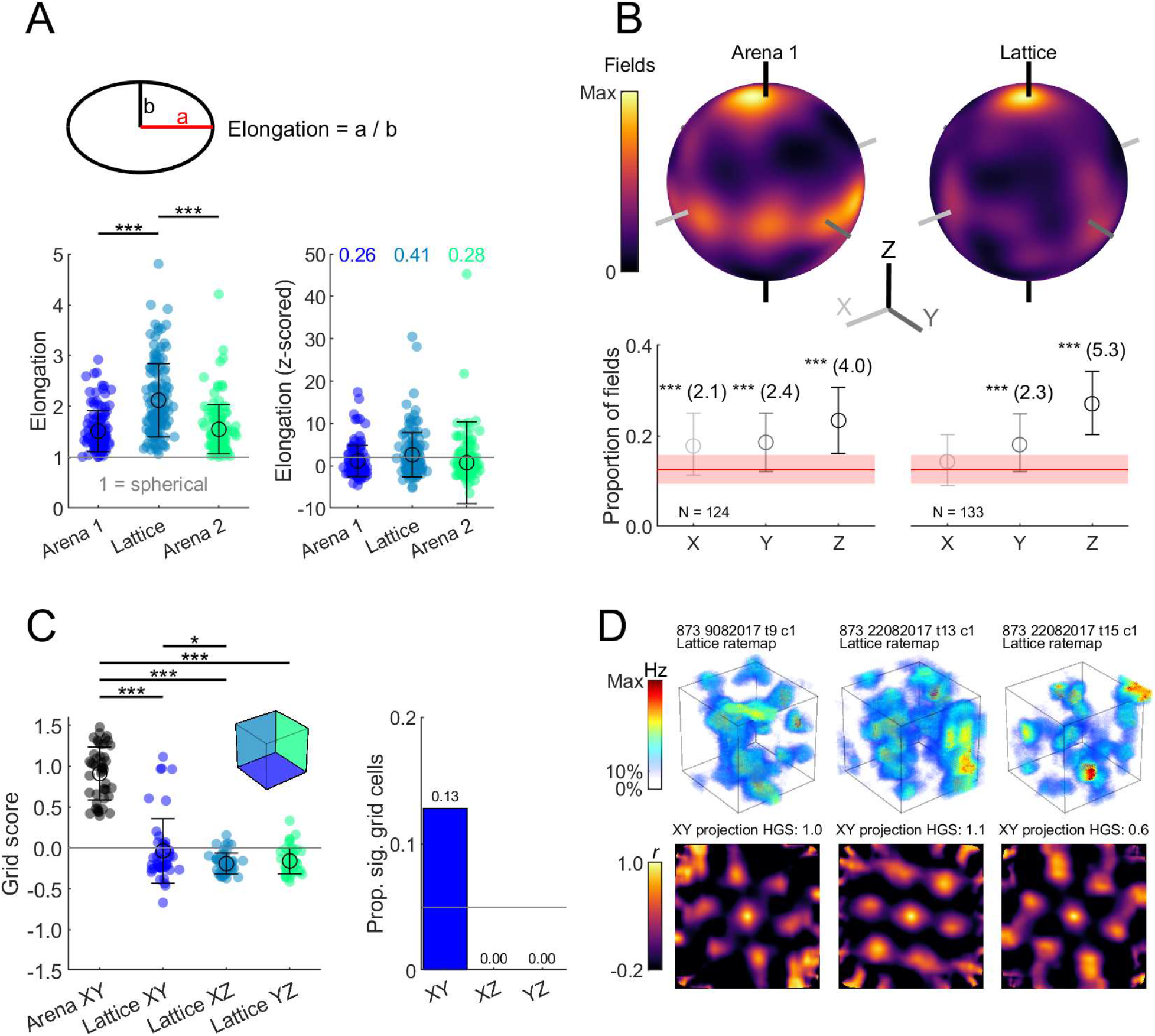
Grid fields in the lattice were vertically elongated and some formed hexagonal columns. (A) left) fields were significantly more elongated in the lattice. Right) Z-scored elongation relative to 100 shuffles; text gives the proportion of fields exceeding the shuffle 95^th^ percentile (i.e., non-spherical; z = 1.96, grey line). (B) Top) spherical heatmaps of the direction of all grid field principal axes (see Fig. S11 for a schematic). Bottom) mean and 95% confidence interval of fields with a principal axis parallel to the X, Y or Z axes relative to chance (red area denotes 99^th^ percentiles). Asterisks denote significant deviation; numbers give effect size (Cohen’s *d*). (C) Left) grid scores for all grid cells in the arena XY plane and each projected plane of the lattice maze. Right) proportion of grid cells with a grid score exceeding the 95^th^ percentile of a chance distribution in each lattice plane. (D) Examples of significant XY grid cells recorded from one rat. Top) volumetric firing rate map. Bottom) autocorrelation of the XY projected firing rate map (HGS: hexagonal grid score).

In summary, we found that rat grid fields did not represent 3D space with a hexagonally close-packed structure (*7, 14*) but instead formed random configurations of slightly vertically-elongated fields. This is consistent with computational models predicting local order in the absence of a regular close-packed structure (*16, 17*). Theta, head-direction, speed coding, spike dynamics and spatial information properties were largely preserved in the lattice maze (Fig. S13-16) suggesting that the lack of grid structure in 3D was not due to a disruption of these inputs. However, field size and spacing increased while speed scores decreased (Fig. S14) and grid spacing modules broke down; findings reminiscent of those seen on a vertical surface (*10*). A failure to integrate distance travelled may explain these results and this may be why place cells were not disrupted in the same maze (*6*): they could use information from other sources such as visual landmarks and borders. This view is supported by the planar hexagonal activity observed in one animal, which could be explained by unusually poor vertical, but not horizontal, integration of distance travelled in this animal (*15, 18*). Alternative explanations for columnar firing fields could include behavioral biases for horizontal movements (*17*) or that overlapping columns represent an efficient way to map higher dimensional space (*19*). This one animal had otherwise no detectable particularity in terms of behavior or recording location.

Together with findings that place cells express spatially-localized firing fields in volumetric space (*6*) and that rats could navigate accurately in the lattice maze (*11, 13*), our results suggest that place cells and spatial mapping can function when grids are irregular. It may be that place cells do not require grid cell inputs for positioning when visual cues are available (*20, 21*). Alternatively, grid cell inputs may not need to be regular to support place cells; the grid symmetry seen in the laboratory may be a consequence of the symmetric geometry and homogeneous behavior of laboratory settings (*22*). These findings thus invite a reappraisal of the computational contributions that grid cells make to spatial mapping, suggesting that any metric contribution of grid fields to spatial localization must arise from the statistics of their dispersal rather than their precise arrangement.

## Acknowledgments

The authors would like to thank Jim Donnett of Axona Ltd. for his expertise in designing and building the experimental equipment.

## Author contributions

Roddy M. Grieves: Methodology, Software, Formal analysis, Investigation, Data Curation, Writing - Original Draft, Writing - Review & Editing, Visualization. Selim Jedidi-Ayoub: Investigation. Karyna Mishchanchuk: Investigation. Anyi Liu: Investigation. Sophie Renaudineau: Methodology. Éléonore Duvelle: Writing - Original Draft, Writing - Review & Editing, Investigation. Kate J. Jeffery: Conceptualization, Resources, Writing - Original Draft, Writing - Review & Editing, Supervision, Project administration, Funding acquisition.

## Data and materials availability

The full raw data set is available from the authors on request. Code to analyze raw data is available from the authors on request.

## Supplementary Materials

Materials and Methods

Figures S1-S16

Tables S1

Movies S1

References (*1-22*)

