## Supplementary Methods and Data for "Grid cell firing fields in a volumetric space"

**This PDF file includes:**

Materials and Methods
Supplementary Text
Figs. S1 to S16
Table S1
Captions for Movies S1

**Other Supplementary Materials for this manuscript include the following:**

Movies S1

### Materials and Methods

Statistics and figures

Unless otherwise stated, we used two-tailed parametric tests (e.g., Matlab *anova1* and *ttest2*) and post-hoc tests compared population means (Matlab *multcompare*, Dunn-Sidak correction). In all figures * = significant at the .05 level, ** = significant at the .01 level, *** = significant at the .001 level. For all dot plots, black lines denote standard deviation, empty circular markers denote the sample mean and filled markers represent individual data points.

Animals

Nine animals were used (only 7 of these animals contributed grid cells), weighing approximately 400–450 g at the start of the experiment. Prior to surgery all animals were housed for a minimum of 8 weeks in a large (2.15m × 1.55m × 2m) cage enclosure, lined on the inside with chicken wire. This was to provide the rats with sufficient 3D climbing experience. During this time, they were given unlimited access to a miniature version of the lattice maze (11cm spacing instead of the 16cm for the recording lattice). Animals were housed individually in cages after surgery and given access to a hanging hammock or climbable nest box for continued three-dimensional experience.

The animals were maintained under a 12 hr light/dark cycle (light starting at 6am) and testing was performed during the light phase of this cycle. Throughout testing, rats were food restricted such that they maintained approximately 90% (and not less than 85%) of their free-feeding weight. This experiment complied with the national [Animals (Scientific Procedures) Act, 1986, United Kingdom] and international [European Communities Council Directive of November 24, 1986 (86/609/EEC)] legislation governing the maintenance of laboratory animals and their use in scientific experiments. A summary of the sessions and cells recorded from each rat can be seen in Table. S1.

Electrodes and surgery

Axona (MDR-xx, Axona, UK) microdrives were used. Drives supported four or eight tetrodes composed of four HML coated, 17 µm diameter, 90% platinum 10% iridium wires (California Fine Wire, Grover Beach, CA) gold plated (Non-Cyanide Gold Plating Solution, Neuralynx, MT) to reduce their impedance to 180–300 kΩ measured at 1Kz. Microdrives were implanted using standard stereotaxic procedures under isoflurane anesthesia (*1*). Briefly, 6 support screws were inserted in the skull, electrodes were implanted after removal of dura, and the drive, skull and screw assembly was secured with an optional first layer of metabond dental cement (Super-Bond C&B Metabond, Parkell, NY, USA) followed by several layers of regular acrylic dental cement (Simplex Rapid Acrylic, Kemdent, Wiltshire, UK). Electrodes were implanted at a posterior-anterior 8-11° slope (with the electrode tip pointing towards the animal's nose), 1-1.5mm below dura, approximately 4.5mm medial to the midline and as close as possible to the transverse sinus. See Fig. S3 for histology results and a photo of an implanted drive.

Apparatus

All experiments were conducted in the same room (3.2×2.1×2.2m) under moderately dimmed light conditions. Three of the room walls were covered with black material with large high-contrast cues on two of them (1.5×1.2m cardboard sheet and a 1×1.7m yellow plastic sheet). The fourth wall was covered with a white cotton sheet. The floor of the room was covered with black anti-static linoleum flooring. We used two pieces of experimental apparatus: a square open field environment (‘arena’) and a cubic lattice composed of horizontal and vertical climbing bars (‘lattice’).

The arena was a 1.2×1.2m square high-walled wooden enclosure, composed of four 1.8×0.65m matte black painted walls (Fig. S1A). The top edges of these walls were covered with large, corrugated tubing to prevent the rats from exploring this area. The bottom edge of this square was highlighted with a strip of 50% grey paint. One 0.45×0.65m matte white wooden cue was affixed to one wall. Rats were recorded freely foraging in the arena for randomly dispersed flavored puffed rice (CocoPops, Kelloggs, Warrington, UK).

The cubic lattice maze (Fig. 1A and B) was constructed from a children's toy-set (Quadro, Hamburg, Germany). Hollow cubes were created by attaching red plastic tubes (length: 150mm, diameter: 10mm) using 6- or 4-way connectors (each 10mm wide). These cubes were then assembled into a 6×6×6 cubic maze (0.97×0.97×0.97m). The maze was raised 0.45m above the ground, initially on black metal stools but later on a narrow wooden frame. To encourage exploration, malt paste (GimCat Malt-Soft Paste, H. von Gimborn GmbH) was affixed to bars of the lattice by the experimenter. This paste was spread evenly throughout the maze, midway along bars, equally between horizontal and vertical bars and reapplied every 15 minutes.

Recording setup and procedure

Single unit activity was collected using a custom built 64-channel recording system (Axona, St. Albans, UK). The rat’s microdrive was connected to a wireless headstage (custom 64-channel, W-series, Triangle Biosystems Int., Durham, NC). Analog signals were transmitted to a wireless base station via dual receiver antennae situated approximately 1m above the maze environments. Unfiltered signals were sampled at 50 kHz, amplified 100 times and transmitted at approximately 3.375 GHz (300 µW at 3m). They were then passed to an Axona pre-amplifier and amplified a further 100 times, then to a system unit for single unit recording where the signal was band-pass (Butterworth) filtered between 300 and 7000 Hz. Signals were digitized at 48 kHz and could be further amplified 10–40 times at the experimenter’s discretion. For LFP recording a 4.8 kHz signal was saved as above which was then band-pass filtered between 6 and 12 Hz ($4^{th}$ order butterworth filter, Matlab *butter* and *filtfilt*) for theta analyses described below. The position of the animal was recorded using five infrared sensitive CCTV cameras (Samsung SCB-5000P) tracking four wide-angle infrared LEDs (Osram Opto SFH 487P, 880nm) fixed to the wireless headstage.

After recovery from surgery, rats were screened for single unit activity and for the presence of theta oscillations once or twice a day, five days a week. Screening was performed in the open field apparatus, after which rats freely foraged on the lattice maze for the same duration. Once the presence of grid cells was confirmed, rats were recorded using the experimental procedure.

In these sessions, rats were recorded for a minimum of 18 minutes in the arena and until they had sufficiently explored the environment (mean ±SD: 27.8 ±3.8 minutes). They were then allowed to rest and drink in an opaque, lidded box for approximately 10 minutes. During this time, the arena was dismantled and replaced with the lattice maze. Rats were then placed on the bottom layer of the lattice and left to forage in this environment for a minimum of 45 minutes and until they had sufficiently explored the environment (mean ±SD: 69.0 ±22.8 minutes). Rats were returned to the opaque box then recorded in the arena for a further minimum of 16 minutes and until they had sufficiently explored the environment (mean ±SD: 27.4 ±9.9 minutes). During recordings, the experimenter monitored progress from a connected room which housed the recording equipment and was separated from the experimental room by a black opaque curtain.

At the end of the recording session, the animals were removed from the apparatus and the electrodes were lowered by at least 20 μm in order to maximize the chance of recording from a different population of cells on the following day. Grid cells with similar fields seen on the same tetrodes on consecutive days were discarded where possible, all analyzed grid cells can be seen in Fig. S2. Rats were tested until grid cells were not observed and it was judged that the electrodes had left the desired layer (mean ±SD: 4.7 ±2.7 sessions).

Trajectory reconstruction

The rat's position was tracked in real time at a 25Hz sampling frequency using DacqTrack software (Axona, St. Albans, UK). Position data were synchronized with neural data using a pulsed optic interface – each camera monitored a 1Hz TTL initiated light source, controlled by the recording system, which allowed accurate, offline synchronization. The onset of these light pulses was used to continually re-align the position data using nearest neighbor interpolation (Matlab *interp1*).

The rat's 3D position was then reconstructed using the direct linear transform algorithm (*2*), applied to the data from all five cameras, in pairs. Briefly, these cameras were first calibrated in order to reverse any distortion introduced by their optical elements (Matlab *estimateCameraParameters*, *undistortImage* and *undistortPoints*). We then imaged the same checkerboard pattern with each camera and used its 3D pose to calculate the distance and orientation of each camera relative to it and thus to each other (Matlab *extrinsics* and *cameraMatrix*). Using this information, we constructed a fundamental matrix. If $x$ are some points viewed by camera 1 and $x'$ are the same points viewed by camera 2, the fundamental matrix, $F$, represents the relationship between points $x$ and $x'$:

$$x_i' F x_i = 0$$

This relationship can be used to triangulate any given pair of points imaged by two cameras into three-dimensional space (*2*). For each recording session we reconstructed the animal's path using every possible pair of cameras (Matlab *triangulate*) and we then combined these reconstructions into one single trajectory. This was achieved by taking the weighted mean of each point, where the weighting was the reliability of the point's estimated location. Reliability was assessed using each point's reprojection error; after triangulation each point was projected back into both camera images; the reprojection error was then calculated as the distance between the original and reprojected position of the point.

In this setup, the rats need only be viewed by two cameras at any one time for a successful reconstruction, allowing for near continuous tracking even in cluttered, complex environments such as the lattice maze. Our cameras were extremely stable; however, re-calibrations were conducted once every two to four weeks to ensure continued reconstruction accuracy. For segments of missing tracking data, we simultaneously interpolated and smoothed the existing data using an unsupervised, robust, discretized, n-dimensional spline smoothing algorithm (Matlab *smoothn* (*3*, *4*)). Example 3D trajectories can be seen in Fig. S1B-C.

Behavior and spherical heat maps

Using smoothed and interpolated 3D reconstructed position data we calculated the instantaneous three-dimensional heading of the animal as the normalized change in position:

$$\hat{u} = \frac{\vec{u}}{||\vec{u}||}$$

where;

$$\vec{u} = (\Delta_X(t), \Delta_Y(t), \Delta_Z(t))$$

and;

$$||\vec{u}|| = \sqrt{\Delta_X(t)^2 + \Delta_Y(t)^2 + \Delta_Z(t)^2}$$

this gives a unit vector representing the animal's heading at time *t*. For visualization we projected these vectors on to a unit sphere as described below (Methods: *Grid field orientation*).

Grobéty and Schenk (*5*) and Jovalekic et al. (*6*) previously reported, in lattice mazes similar to the one used here, that rats exhibited a strong bias for horizontal movements. To test this in our lattice maze, we compared the number of times each animal crossed from one lattice element (the smallest cubic subcomponent of the maze) to another. This was computed in each of the $X$, $Y$ and $Z$ axes as the lattice edges and bars were aligned with these. We considered a 'crossing' to have occurred when the head of the rat moved from one unit to another. A detailed analysis of the behavioral data can be found elsewhere (*7*) and so only brief results related to grid cell activity are reported here.

#### Spike sorting

Single-unit activity was analyzed offline using a combination of Matlab functions and spike-sorting software. First, the dimensionality of the waveform information was reduced to the first three principal components and amplitude. Based on these parameters, an automated spike sorting algorithm (Klustakwik v3.0, (*8*)) was used to distinguish and isolate separate clusters. The clusters were then further checked and refined manually using a cluster cutting GUI (TINT v4.4.12, Axona, UK). As well as the previously mentioned features, manual cluster cutting also made use of spike auto- and cross-correlograms.

#### Firing rate maps

To generate three-dimensional volumetric firing rate maps we used an adaptive binning method described previously (*11*, *12*). Briefly, for every bin a circle centered on the point was gradually expanded until the following criterion was met:

$$r > \alpha / n\sqrt{s}$$

where $\alpha$ is a constant, $r$ is the radius of the circle in pixels, $n$ is the number of occupancy samples falling within the circle, and $s$ is the total number of spikes falling within the circle. Once this criterion was met, the firing rate assigned to the point was equal to $s/n$. For our maps, $\alpha$ was set to the value 1600 and we calculated firing rate across a 2.5cm$^3$ grid. Periods where the animals were moving <5cm/s were not included in any firing rate maps to avoid potential contamination by low-speed phenomenon (sharp-wave/ripple activity, licking behavior etc.).

When calculating the stability of spatial activity between volumetric maps to avoid inflated comparisons based on many small voxels, we instead made maps using a standard histogram procedure (Matlab *histcn*, B. Luong). For this, data were binned using a much larger 10cm$^3$ grid and smoothed using a Gaussian kernel with a standard deviation of 2.5 bins (Matlab *imgaussfilt3*).

For two-dimensional maps, such as the Cartesian planar projections, we generated firing rate maps using a standard histogram procedure (Matlab *histcounts2*). For this data were binned using a 2.5cm$^3$ grid and smoothed using a Gaussian kernel with a standard deviation of 1 bin (Matlab *imgaussfilt*).

Recording stability between arenas

To verify that cells were stably recorded during our maze sessions we used a similar approach to one described previously (*13*); first we computed the Pearson pairwise correlation (Matlab *corr*) between the first and second arena ratemaps recorded before and after each lattice maze session. For this we used the two-dimensional XY projected ratemaps described above (Methods: *Firing rate maps*). For comparison we correlated first and second arena sessions from random grid cells whilst maintaining their temporal order (i.e., first arena vs second arena from a random cell). This shuffle was repeated 5000 times. The results of this analysis can be seen in Fig. S4.

Spatial stability within sessions

To test the within-session stability of spatial representations we divided maze sessions into two halves of equal length (first 50% and second 50%) and computed the Pearson pairwise correlation (Matlab *corr*) between the firing rate maps for these halves. For recording stability, two-dimensional maps were generated from data projected onto the Cartesian coordinate planes. However, we also compared volumetric maps generated as multivariate histograms with $10cm^3$ voxels smoothed using a Gaussian with a 2.5 bin standard deviation (Matlab *imgaussfilt3*). Large bins and smoothing were used here to reduce the huge number of spatial bins compared between maps (from around the $9\times10^4$ voxels found in adaptively binned ratemaps to around $2\times10^3$ voxels). Examples of these maps can be seen in Fig. S5A. In both cases we compared the observed correlation values to shuffled distributions generated by comparing session halves from random cells (i.e., first 50% vs second 50% from a random cell). This shuffle was repeated 5000 times for each map type. The results of this analysis can be seen in Fig. S5.

Spatial information and sparsity shuffles

To determine if the firing of grid cells was less homogeneous (i.e., more 'clumped') than chance we generated firing rate maps using a standard histogram procedure (Matlab *histcn*, B. Luong) with $2cm^3$ voxels and smoothed using a Gaussian kernel with a standard deviation of 2 voxels (Matlab *imgaussfilt3*). We then calculated spatial information content (bits/second) as:

$$spatial\ information = \sum_{i=1}^{N} p_i \frac{\lambda_i}{\lambda} log_2 \frac{\lambda_i}{\lambda}$$

Next, for each grid cell we shuffled its spike train 100 times by random increments of 0.02s (minimum 20s) and for each shuffle recomputed a firing rate map and spatial information as above. Lastly, we expressed the observed spatial information values in standard deviations from the shuffle:

$$standardised\ value = \frac{(observed\ value - \mu_{shuff})}{\sigma_{shuff}}$$

This essentially z-scores the observed values relative to the shuffles. Scores greater than 1.96 exceed the shuffle 95$^{th}$ percentile and thus deviate significantly from the shuffle at the .05 level. Similar results were obtained using sparsity (data not shown).

Directional analyses and shuffles

As we did not have access to 3D head direction information, we estimated instantaneous projected azimuthal head direction as:

$$\theta(t) = tan^{-1}[\Delta y^t / \Delta x^t]$$

where $\Delta x^t$ and $\Delta y^t$ represent the change in X or Y position respectively (*14*) for which we used the smoothed and interpolated position tracking (Methods: *Trajectory reconstruction*) ignoring movements in the Z axis. To generate head direction tuning curves, we binned position and spike directions into 6° bins, smoothed the resulting histograms with a Gaussian kernel (Matlab *imgaussfilt*, σ = 3 bins, circular padding) and then generated a tuning curve by dividing the spike histogram by the time spent in each bin. From these we calculated, as a measure of directionality, the Rayleigh vector length (Matlab *circ_r*, circular statistics toolbox (*15*)) and preferred firing direction (PFD) as the bin containing the maximum firing rate.

Next, for each cell we shuffled its spike train 100 times by random increments of 0.02s (minimum 20s) and for each shuffle recomputed a directional rate map and statistics as above. A cell was categorized as directionally modulated if it exhibited a Rayleigh vector greater than the 95th percentile of the shuffled values in both arena sessions and fired at a rate greater than 0.1Hz in both.

Lastly, to determine if directionally modulated cells maintained their allocentric firing directions across mazes (all recordings were made in the same location in the same room) we correlated the tuning curves for all cells as a population in the arena to their tuning curves in the lattice. For comparison we circularly shifted each cell's lattice tuning curve independently by a random number of bins (between 1 and 60) and recomputed the correlation. We repeated this 1000 times and if the original correlation exceeded the 95th percentile of the shuffled values we considered that the cell popuation firing was more stable than chance; this difference was also expressed as a standardized Z-score as described above (Methods: *Spatial information and sparsity shuffles*).

Autocorrelations

Two- and three-dimensional autocorrelations, $r$, of each grid cell's firing rate map were calculated according to:

$$r(\tau_x, \tau_y, \tau_z) = \frac{M \sum_{x,y,z} \lambda(x,y,z)\lambda(x-\tau_x, y-\tau_y, z-\tau_z) - \sum_{x,y,z} \lambda(x,y,z) \sum_{x,y,z} \lambda(x-\tau_x, y-\tau_y, z-\tau_z)}{\sqrt{\left[M \sum_{x,y,z} \lambda(x,y,z)^2 - \left[\sum_{x,y,z} \lambda(x,y,z)\right]^2\right] \left[M \sum_{x,y,z} \lambda(x-\tau_x, y-\tau_y, z-\tau_z)^2 - \left[\lambda(x-\tau_x, y-\tau_y, z-\tau_z)\right]^2\right]}}$$

where $\lambda(x, y, z)$ is the firing rate at the location $(x, y, z)$ in the firing rate map, $M$ is the total number of voxels in the rate map, and $\tau_x$, $\tau_y$ and $\tau_z$ correspond to $x$, $y$, and $z$ coordinate spatial lags (*14*).

Autocorrelation self-similarity

As a measure of firing rate map self-similarity along each axis we extracted, for each grid cell, the autocorrelation values falling along the autocorrelation midlines (Matlab *interp3* with linear interpolation). Intuitively, if a volumetric firing rate map contained circular columns that spanned the entire z-axis then the autocorrelation would also contain a columnar peak in its center, spanning the entire z-axis. Thus, values falling along a line drawn straight down the middle of the autocorrelation from top to bottom would all be high, while values falling on a line drawn straight across the autocorrelation from one side to the other through the middle would peak near the center but otherwise contain low values.

Autocorrelation correction for anisotropy

As in place cells (*16*) some grid fields were significantly elongated, if these fields formed a close-packed arrangement this could take two forms. First, fields could be arranged isotropically (field centroids forming equilateral tetrahedra) while remaining individually elongated, perhaps resulting in the overlap of vertical layers. While this should not present a problem for the planar symmetry analysis proposed by Stella and Treves (*17*) a second possibility is that the underlying arrangement itself could be elongated (field centroids forming acute and obtuse tetrahedra) with fields elongating to fill the volume between them. This latter configuration would disrupt the angular relationships between packing layers and the planar symmetry analysis.

To account for this, we repeated the planar analysis after correcting grid cell autocorrelograms to remove anisotropy introduced by field elongation. This would also correct anisotropic close-packed arrangements allowing FCC and HCP arrangements to be identified. For this, we thresholded autocorrelations at a correlation value of 0.25 (Matlab *imbinarize*) and extracted the central region. This central region should reflect the average characteristics of the firing fields in the firing rate map, their elongation and orientation. Using this fact, we resized the autocorrelation along each dimension by the amount necessary to 'correct' this central peak into a sphere (Matlab *imresize3* with cubic interpolation and antialiasing); thus, correcting any anisotropy in a potential close-packed arrangement (see Fig. S10A for an example).

Grid score

Gridness scores were calculated similarly to prior papers (*18*, *19*). The 2D autocorrelogram was thresholded to leave only values >0.3 and the seven most central correlation peaks were found. The peak closest to the center of the autocorrelation was excluded and the annulus concentric with the autocorrelogram that contained the other six peaks was isolated. The inner/outer radii defining this annulus were chosen as $\pm r$, where $r$ was the estimated radius of the most central peak. Pearson correlations between rotationally offset copies of the annulus were computed. Gridness score (also called hexagonal gridness score, HGS) was calculated as the minimum correlation obtained at rotational offsets 60° and 120° minus the maximum obtained at 30°, 90°, and 150° which results in a high value when the autocorrelation exhibits a hexagonal structure and a low value otherwise. In addition we calculated an equivalent score for a square firing pattern (also called square gridness score, SGS) as the minimum correlation obtained at rotational offsets 90° and 180° minus the maximum obtained at 45°, 135°, and 225° (*14*).

Grid score shuffles

To determine if a cell's hexagonal grid score or other spatial parameter was greater than could be expected by chance, for each session we used a bootstrap vs shuffle approach (*20*, *21*). First, estimated spatial parameter values (HGS, SGS, speed score) were obtained by resampling spikes using a bootstrap with replacement procedure (100 iterations). At each iteration we recreated a firing rate map and spatial autocorrelation and recalculated the spatial parameter values. Spatial parameter values were then estimated as the median of the collected bootstrapped values.

Next, we repeated the same procedure (100 iterations) but instead of resampling spikes we circularly shifted the spike train of the cell by a random 0.2s increment (greater than 20 seconds). If the median parameter value obtained from bootstrapping exceeded the 95$^{th}$

percentile of the values obtained from the shuffles, the spatial parameter was considered to be greater than expected by chance.

Grid cell criteria

After completing the parameter shuffles described above a cell was considered a grid cell if its grid score exceeded that expected by chance in both arena sessions (recorded before and after the lattice session). The results of this analysis can be seen in Fig. S4B to D.

Grid fields

Firing fields were detected in adaptive binned firing rate maps (Methods: *Firing rate maps*) as regions of more than 64 contiguous voxels with a firing rate greater than 30% of the map's peak value (Matlab *imbinarize* and *regionprops3*). Additionally, each field had to have a peak firing rate greater than 1Hz and rats had to visit it more than 5 times during a session. We then extracted field properties such as volume, centroid, principal axis lengths, eigenvectors and eigenvalues using previously established methods (*16*)(Matlab *regionprops3*).

Grid fields per cubic meter

To estimate the practical volume of our 3D mazes we calculated an average dwell time map across all sessions and animals. We then thresholded this map so that only bins containing an average dwell time >0.1s remained. Maze volume was then estimated as the total volume of these remaining voxels. In this way the arena was estimated as $0.4m^3$ and the lattice as $1.2m^3$.

Grid field size

Grid field size was estimated as the radius of a sphere with a volume equivalent to that of the 3D autocorrelation central peak after thresholding at 0.25. This method was validated on simulated FCC arrangements with different field sizes (Fig. S6A&B).

Grid field spacing

Grid spacing was estimated by expanding a sphere outward from the center of the 3D autocorrelation, in steps of one bin up to a radius 0.6 times the maximum side length of the autocorrelation, and calculating the median correlation in bins within a distance of 2 bins from the surface of the sphere. Average field spacing was then estimated as the location of the first peak in the median correlation values after excluding the central autocorrelation peak (Matlab *findpeaks*, with a minimum peak prominence of 0.01 and excluding peaks at a distance <5 bins). This method was validated on simulated field arrangements, and correctly estimates grid spacing regardless of the underlying configuration (Fig. S6C&D). The average median correlation values for all grid cells can be also be seen in Fig. S6E.

Grid field orientation

We extracted each grid field's orientation and principal axes, which were defined as the orientation and major axes of an ellipsoid with the same normalized second central moments as the field region. In more detail, we calculated the second central moments or covariance matrix which best described a thresholded place field, in effect fitting a multivariate normal distribution to the field. The direction and magnitude of the best-fit ellipse which describes the place field are then given by the eigenvectors and eigenvalues of this covariance matrix respectively (Matlab *regionprops3*).

To determine if fields were oriented in three dimensions along one or more arbitrary axes we projected the field eigenvectors and their antipodal equivalents onto a unit sphere. We then extracted the number of fields falling within regions on the surface of the sphere corresponding to the intersection of the sphere and the Cartesian XYZ axes. These regions are equivalent to ~60° conic sections centered on each respective axis in one direction from the origin, thus for each axis we combined the two corresponding directional regions.

To determine whether more fields were parallel to an axis than would be expected by chance, we generated 1000 random points on the face of a sphere and counted the proportion of points falling within the area around each axis. We did this 1000 times. Chance was calculated as the interval between the $2.5^{th}$ and $97.5^{th}$ percentile ranks of this distribution. If the observed field count for an axis exceeded the upper threshold it was considered to be overrepresented with respect to chance.

For visualization, we calculated the Von Mises–Fisher kernel smoothed density estimate of these grid field vectors across the sphere's surface. Briefly, the Gaussian used was defined as:

$$g(x) = e^{\left(-0.5\left(\frac{x}{\sigma}\right)^2\right)}$$

where $x$ was defined as the inverse cosine of the inner dot product between each vector point and points across a sphere's surface (Matlab *sphere*) and $\sigma$ was the standard deviation of the Gaussian, which was set to 10. In this way, the resulting three-dimensional heat plots give a density estimate of points on the sphere, where density is estimated as the sum of the Gaussian weighted distances (along the surface of the sphere) to every data point. These 3D spherical maps are presented in the main text for visualization only. See Fig. S11 for a schematic explanation.

Grid field elongation

After extracting a field's principal axis lengths (Methods: *Grid field orientation*) we calculated the elongation index as:

$$Elongation = \frac{P1}{0.5(P2 + P3)}$$

where $P1$, $P2$ and $P3$ are the principal axes from largest to smallest, respectively. This gives a measure of the curvature of the place field: large elongation values represent elongated fields while a value of 1 would represent a sphere. In the case of the arena, elongation was calculated using the first two principal axes ($P1/P2$).

We then tested whether this elongation index deviated significantly from a distribution that would be expected by chance using an analysis inspired by one reported previously (*22*). For each field we defined a perfect sphere, centered on the field's centroid. The diameter of this sphere was calculated such that it would share the same convex volume (the volume of the convex hull enclosing the field voxels) as the place field. This was calculated as:

$$equivalent\ diameter = 6\left(\frac{vf}{\pi}\right)^{\frac{1}{3}}$$

where *vf* is the convex volume of the place field. This is more accurate than the geometric mean approach reported previously (*22*) which assumes all place fields are perfectly elliptical and thus tends to underestimate equivalent diameter. Next, the spikes emitted within the place field were randomly shuffled among the trajectories through this sphere using a multivariate Gaussian process (Matlab *normrnd*). The mean of the Gaussian was the sphere center, and the standard deviation of the Gaussian was set to 1.8 × the radius of the sphere (to approximate the 20% thresholding used during field detection). Each spike was then assigned to the position of the nearest trajectory data point (Matlab *knnsearch*). The result of this procedure was a normally distributed point cloud of spikes centered on the centroid of the original field with the same equivalent diameter and firing rate.

We then recomputed the firing rate map for these shuffled spikes and extracted its elongation index as described above. This procedure was repeated 100 times for each place-field. Place fields with an elongation index that could be expected, by chance, from an underlying spherical field (i.e., with an elongation index lower than the 95$^{th}$ percentile rank of the shuffled distribution) were defined as spherical or isotropic: otherwise, place fields were defined as non-spherical, anisotropic, or elongated.

Planar symmetry analysis

Face-centered-cubic (FCC) is a cubic lattice structure that results from stacking hexagonally arranged layers of spheres in the sequence ABC, where layers B and C are offset hexagonal patterns that rest in the spaces between the spheres below. Hexagonal close-packed (HCP) is a similar lattice structure that describes an ABA sequence (Fig. 1A, Fig. S7A)(*14*, *17*, *23*, *24*).

Where the autocorrelogram of a 2D grid cell firing rate map resembles a hexagonal grid, centered on a central peak, the autocorrelogram of a 3D FCC configuration also reproduces the same 3D pattern spanning around a central peak (*17*). If the field configuration is aligned to the XY axes, a horizontal plane cut through the center of the autocorrelation will pass through a hexagonal field arrangement and the correlation values falling on this plane will present a high grid score. Additionally, three further planes angled at a pitch of 72° from the XY plane can be found that also pass-through hexagonal arrangements and present high grid scores. These planes will also be arranged with 120° between them in azimuth. Additionally, due to the cubic nature of the FCC arrangement (another name for face-centered cubic is cubic-close-packed), there are also 3 planes at a pitch of 57° from the XY plane arranged with 120° between them in azimuth, offset from the 72° hexagonal planes by 60° in azimuth, that transect fields arranged in a square formation. In this way, extracting every possible plane through the autocorrelation center, and mapping their hexagonal and square grid scores (HGS and SGS respectively) in turn, can be used to determine the likelihood of an FCC arrangement (Fig. S7B-C, Fig. S8).

By contrast the autocorrelation of an HCP configuration, while still centered around a peak, does not exactly resemble the original pattern. While there is still a 'best plane' corresponding to a horizontal shift of the grid pattern (like in the 2D case) due to the half overlap of peaks between layers there are a further six layers that pass-through peaks resembling a hexagonal grid (*17*). Assuming a field arrangement aligned to the XY axes a horizontal plane cut through the center of the autocorrelation will pass through a hexagonal field arrangement and the correlation values falling on this plane will present a high HGS. Additionally, six further planes angled at a pitch of 72° from the XY plane can be found that also pass-through hexagonal arrangements. However, three of these planes present high HGS (arranged with 120° between them in azimuth) while the other three present low HGS (arranged 60° offset to the other planes in azimuth). In an HCP arrangement planes with square field arrangements can also be found, again at 57° from the XY plane in pitch and at the same azimuthal angles as the hexagonal ones (Fig. S7C, Fig. S8).

Last, the autocorrelation of a hexagonal columnar field arrangement is the same arrangement centered around a columnar peak. Assuming the columns are parallel to the z-axis we would expect horizontal slices to present high grid scores which would decrease as the pitch of the slices diverges from the horizontal (Fig. S7C, Fig. S8).

In these examples we assumed the configurations are aligned to the XY axes, but once grid scores are mapped for every azimuth and pitch slice combination it is instead possible to estimate the 3D orientation of the arrangement using the 'best plane' or the transecting plane with the highest grid score. In the FCC case all four planes are equally 'best,' but this is useful for HCP and columnar arrangements. In all cases it allows us to correct grid cell autocorrelations, rotating them so that the best plane is always horizontal (Fig. S9A).

Structure scores ($\chi_{CP}$, $\chi_{FCC}$, $\chi_{HCP}$ & $\chi_{COL}$)

Once we collected the HGS and SGS for every possible plane transecting a grid cell autocorrelation we looked to calculate scores that could be used to differentiate the different field configurations. We used an approach similar to that proposed by Stella and Treves (*17*) but also took into account the square grid scores which allowed us to differentiate FCC and HCP arrangements based solely on their autocorrelations. We also extended this analysis to include a score for columnar arrangements (Fig. S8).

We built volumetric firing rate maps (Methods: *Firing rate maps*) and autocorrelations (Methods: *Autocorrelations*) and then extracted planes transecting the autocorrelation (65 pitch angles and 65 azimuth angles for a total of 4225 planes; Matlab *sphere* and *obliqueslice*). For each plane we calculated its HGS and SGS (Methods: *Grid score*). We next found the 'best plane' as the one associated with the maximum HGS and corrected the autocorrelogram so that this plane would form the XY horizontal (Fig. S9A). We then interpolated the HGS and SGS maps up to 129 pitch and azimuth angles using a spherical nearest neighbor method where nearest neighbors were found based on the inverse cosine of the inner dot product between each vector point and points across a sphere's surface (Matlab *sphere*).

Next, we extracted the HGSs found at a 72° pitch from the best plane and the SGSs found at a 50° pitch from the best plane. As both FCC and HCP are expected to exhibit high HGSs and SGSs at these respective angles we calculated a general quality score ($\chi_{CP}$) as the median of these two distributions combined.

For an FCC score ($\chi_{FCC}$) we found the three 120° offset azimuth angles at 72° pitch from the best plane with the maximum total hexagonal grid score. We then calculated $\beta$ as the median square grid score found at the same azimuthal angles and at a 50° pitch and $\alpha$ as the median square grid score found at 60° offsets from these in azimuth and at a 50° pitch (Fig. S8 left). From these:

$$\chi_{FCC} = \alpha - \beta$$

For the HCP score ($\chi_{HCP}$) we found the three 120° offset azimuth angles at 72° pitch from the best plane with the maximum total hexagonal grid score. We then calculated $\alpha$ as the median square grid score found at the same azimuthal angles and at a 50° pitch and $\beta$ as the median square grid score found at the same azimuthal angles and at a 72° pitch (Fig. S8 middle). From these:

$$\chi_{HCP} = \alpha - \beta$$

For the columnar score ($\chi_{COL}$) we calculated $\alpha$ as the median hexagonal grid score found at all pitch angles within 60° of the best plane and $\beta$ as the median of all remaining grid scores (Fig. S8 right). From these:

$$\chi_{COL} = \alpha - \beta$$

Simulated field arrangements

Where $S$ was the side length of a hexagon, points were separated by $S$ along the x axis and $\sqrt{3}(0.5S)$ along the y axis with every second row of points along the y axis offset by $S/2$ in x; this gives points tiling horizontal space in a hexagonal arrangement. For our simulations $S$ was a value randomly drawn for each cell from the uniform distribution between 200 and 600mm. For close-packed arrangements these horizontal points were then stacked along the z axis separated by $(\sqrt{6}/3)S$. For HCP, every second layer along the z axis (2nd onwards) was offset in y by $\sqrt{3}(1/3)\,S$. For FCC, every third layer (2nd onwards) along the z axis was offset in y by $\sqrt{3}(1/3)\,S$ and every third layer (3rd onwards) along the z axis was offset in y by $-2(\sqrt{3}(1/6)S)$. For a columnar configuration, the horizontal hexagonal points were simply stacked continuously in z. For a random arrangement of fields, we generated *N* random points (Matlab *rand*), where *N* was the number of points generated for an HCP arrangement, spread within the cuboid volume occupied by an HCP arrangement.

When the desired arrangement of points was generated, for every pixel of the simulated ratemap we calculated the Euclidean distance to the nearest field point (Matlab *bwdist*) and weighted these using the Gaussian:

$$g(x) = e^{\left(-0.5\left(\frac{x}{\sigma}\right)^2\right)}$$

where $x$ is the Euclidean distance and $\sigma$ is the Gaussian standard deviation which was set to 2. To ensure that the planar symmetry analysis was capable of differentiating field arrangements even when they are not aligned to the maze/gravity, the resulting simulated firing ratemap was then rotated 30° around a random 3D axis (Matlab *imrotate3* and *rand*). Simulated field arrangements without this last rotation step can be seen in (Fig. S7A).

Grid field shuffle

To generate fields in random positions while remaining as close to the real data as possible we employed a field shuffling technique described previously (*25*). Briefly, adaptive binned firing rate maps were over-smoothed (Methods: *Firing rate maps*, Matlab *imgaussfilt3*, sigma 3 bins) and firing fields were segmented through a watershedding procedure. For this, field peaks were detected as local maxima (Matlab *imextendedmax* with a H-maxima of 0.2) and watersheds were calculated on the distance transform (Matlab *bwdist*) of these peaks (Matlab *watershed*).

The following analyses were then performed on the unsmoothed adaptive binned firing rate map: $N$ uniformly random points were identified in an empty copy of the firing rate map where $N$ was the total number of fields identified in the watershed procedure. For each segmented field (numbered 1 to $N$) the bin with the peak firing rate was copied to one of these random positions. Next, firing rate values were iteratively moved from the original rate map (O) to the new empty copy (E) following this procedure: for each field in turn (1 to $N$) the bin closest to the peak was moved from O to E whilst maintaining its position relative to the peak as closely as possible (the 'ideal' position). Values were not moved to the locations of unvisited bins in O nor could they overwrite values already moved to E, so where the ideal position was not available values were instead moved to the position nearest to it in cityblock distance. This procedure was repeated for every field in turn and in the field order 1 to $N$ iteratively until every bin had been moved from O to a position in E.

Intuitively, this method randomly shuffles the positions of fields within a firing rate map but also preserves, as much as possible, the internal structure of each field. Furthermore, because every firing rate value is moved and both maps share the same number of elements both maps have the same distribution of firing rate values, the same peak firing rate and the same number of unvisited bins. Unvisited bins also retain their spatial positions which maintains the shape and configuration of any uneven sampling in the original firing rate map. Due to the computational time cost associated with this shuffling method when using three dimensional maps and the already large time cost of the planar symmetry analysis, this shuffle was only performed twice per cell. Example field shuffled firing rate maps can be seen in (Fig. S10C).

Local field potential (LFP) analyses

Before analysis, all LFP data were removed of their direct current offsets, slowly changing components, and running line noise using the Chronux toolbox (*26*) *locdetrend* function which subtracts the linear regression line fit within a 1s moving window. They were then resampled at 250 Hz using a polyphase anti-aliasing filter (MATLAB function *resample*, pchip interpolation).

To obtain a theta phase angle for each spike, LFPs were first bandpass filtered in the 6-12 Hz range (fourth-order Butterworth, Matlab *butter* and *filtfilt*) before a Hilbert transform was applied to obtain the instantaneous phase angle (Matlab *hilbert*). Instantaneous frequency was calculated as the derivative of this analytic signal (Matlab *instfreq*) and instantaneous amplitude was calculated as its magnitude.

To assess the relationship between running speed and the theta oscillation we compared the instantaneous theta power/amplitude at every position data point (every 20 ms) to the animals' instantaneous running speed. Instantaneous speed was estimated as the total distance travelled in every 40 ms window. To quantify the relationship between speed and power we fitted a linear regression model using a least-squares approach (Matlab *polyfit*, 1 degree) and extracted the slope, y-intercept and sum of squared error. We also performed the same procedures to test the relationship between running speed and instantaneous frequency.

To calculate general theta characteristics, we computed average power spectral densities (PSDs) for each recording session by first zero-padding LFP data to the next highest power of 2. A Welch spectral estimator was then applied to obtain the PSD (Matlab *pwelch*, Hamming window, 8 segments, 50% overlap). This was computed for 500 logarithmically spaced points between 0-250 Hz. Theta power was estimated as the maximum power found in the theta band (6-12Hz), theta frequency was defined as the frequency associated with this maximum power.

Running speed analyses

Instantaneous running speed was estimated as the total distance travelled in every 40ms window. For each cell, instantaneous firing rate was estimated as the smoothed spike histogram (20 ms bins, 13 bin or 260 ms Gaussian smoothing window using Matlab function *fspecial*). To quantify the relationship between speed and firing rate we used an analysis similar to that described previously (*27*). We binned the animals' running speeds in 2cm/s increments and calculated the mean firing rate for each running speed bin and the total time spent moving at that speed. We then fitted a linear regression model to the average firing rate/speed data using a least-squares approach (Matlab function *polyfit*, 1 degree) and extracted the slope, y-intercept and sum of squared error.

Spike phase and autocorrelation analyses

To quantify the intrinsic theta modulation of every place cell we used an analysis described previously (*28*, *29*). For each cell we calculated the ±500ms spike autocorrelation in 10ms bins, normalized this to the maximum value found between 100 and 150 ms and removed values >1. Then we fit the following function to the remaining data:

$$y(t) = \left(a * \left(\sin\left(2\pi\omega t + \frac{\pi}{2}\right) + 1\right) + b\right) * \exp\left(-\frac{|t|}{\tau_1}\right) + c * \exp\left(-\frac{r^2}{\tau_2^2}\right)$$

where $a, b, c, \omega, \tau_1$ and $\tau_2$ were fit to the data using a non-linear least squares method (Matlab *fit*) and $t$ is the autocorrelogram time lag. In simple terms this function fits a sine wave of frequency $\omega$ to the data and the exponential term allows for this to decrease exponentially as the time lag increases (reflecting the exponential decay inherent to all spike autocorrelations). The last Gaussian term helps to center the fit on the autocorrelogram peak, which we found to be unnecessary in most cases. A measure of theta modulation strength was defined as $a/b$, which intuitively corresponds to the ratio of the sine fit relative to the baseline in the autocorrelogram. The parameter $\omega$ was extracted as the intrinsic theta modulation of the cell. We restricted possible values for $\omega$ to [6, 12], $a$ and $b$ were restricted to non-negative values [0 Inf], $c$ was restricted to [0, 0.8], $\tau_1$ was unrestricted and $\tau_2$ was restricted to [0, 0.05]. This fitting procedure was only carried out on cells that fired at least 500 spikes.

For each cell, the instantaneous theta phase of every spike was calculated by linear interpolation of the instantaneous theta phase signal described previously. These phase angles were binned between $-\pi$ and $\pi$ in 0.1 rad bins. The cell's preferred theta phase was defined as the circular mean of these angles and the strength of this modulation was defined as the mean resultant vector length of these angles (Matlab *circ_mean* and *circ_r* respectively, circular statistics toolbox, (*15*)).

Histology

At the end of the experiment animals were given an overdose of pentobarbital intraperitoneally (Euthatal, Merial Animal Health Ltd., Essex, UK), and perfused with 0.9% saline solution followed by a 4% formalin solution. The brain was extracted and stored in 4% formalin for at least seven days prior to any histological analyses. The brains were sliced sagittally in 30 µm sections on a freezing microtome at −20°. These sections were stained with a 0.1% cresyl violet solution and the slice best representing the electrode track was then imaged. Histology results for every animal can be seen in Fig. S3.

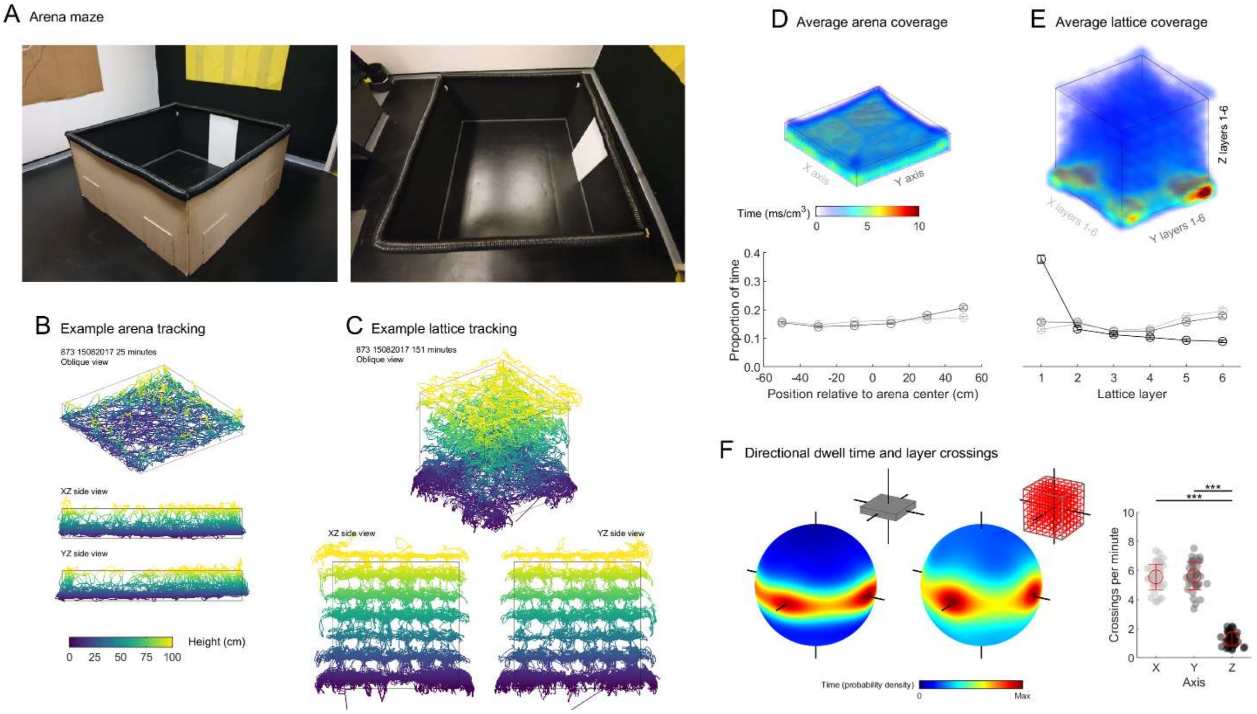


**Fig. S1. Animals explored the whole lattice maze with a bias for horizontal movements.** (A) Side and top-down views of the arena. (B) Different views of the 3D positions for a representative arena session. (C) Different views of the 3D positions for a representative lattice maze session. (D) Top) averaged dwell time histogram for all arena sessions; Bottom) mean ± SEM proportion of time spent at different positions within the arena averaged across sessions. Rats explored the entire arena homogenously. (E) Top) averaged dwell time histogram for all lattice maze sessions; bottom) Mean ± SEM proportion of time spent in each layer of the maze, averaged across sessions. Rats explored the entire lattice homogenously in X and Y but with a strong vertical bias for the bottom layer. (F) Left) spherical heatmaps showing the time spent moving at every possible yaw × pitch angle in the arena and lattice. Inset schematics give the maze shape and the corresponding axes shown extending through the spheres. Right) Markers represent sessions, as reported previously (*7*, *16*), rats were biased towards horizontal movements in both mazes and were far more likely to move parallel to the maze axes: the walls of the arena or the sides and bars of the lattice ($F(2,123) = 417.2$, $p < .0001$, $\eta^2 = 0.872$, Z vs X or Y $p < .0001$, all other $p > .05$, one-way ANOVA).

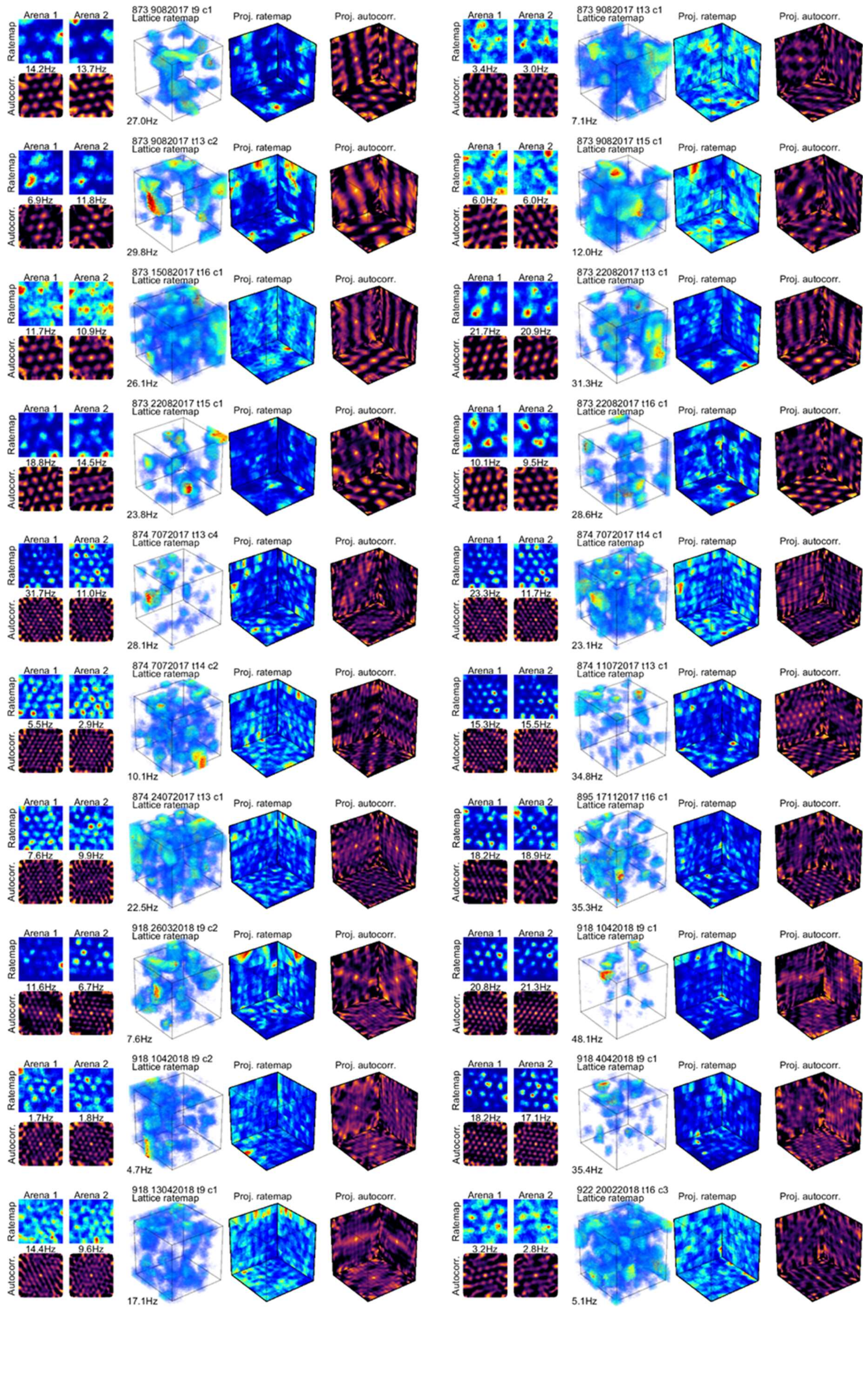
Arena 1
Arena 2
Ratemap
Autocorr.
14.2Hz
13.7Hz
873 9082017 t9 c1
Lattice ratemap
Proj. ratemap
Proj. autocorr.
27.0Hz
Arena 1
Arena 2
Ratemap
Autocorr.
3.4Hz
3.0Hz
873 9082017 t13 c1
Lattice ratemap
Proj. ratemap
Proj. autocorr.
7.1Hz
Arena 1
Arena 2
Ratemap
Autocorr.
6.9Hz
11.8Hz
873 9082017 t13 c2
Lattice ratemap
Proj. ratemap
Proj. autocorr.
29.8Hz
Arena 1
Arena 2
Ratemap
Autocorr.
6.0Hz
6.0Hz
873 9082017 t15 c1
Lattice ratemap
Proj. ratemap
Proj. autocorr.
12.0Hz
Arena 1
Arena 2
Ratemap
Autocorr.
11.7Hz
10.9Hz
873 15082017 t16 c1
Lattice ratemap
Proj. ratemap
Proj. autocorr.
26.1Hz
Arena 1
Arena 2
Ratemap
Autocorr.
21.7Hz
20.9Hz
873 22082017 t13 c1
Lattice ratemap
Proj. ratemap
Proj. autocorr.
31.3Hz
Arena 1
Arena 2
Ratemap
Autocorr.
18.8Hz
14.5Hz
873 22082017 t15 c1
Lattice ratemap
Proj. ratemap
Proj. autocorr.
23.8Hz
Arena 1
Arena 2
Ratemap
Autocorr.
10.1Hz
9.5Hz
873 22082017 t16 c1
Lattice ratemap
Proj. ratemap
Proj. autocorr.
28.6Hz
Arena 1
Arena 2
Ratemap
Autocorr.
31.7Hz
11.0Hz
874 7072017 t13 c4
Lattice ratemap
Proj. ratemap
Proj. autocorr.
28.1Hz
Arena 1
Arena 2
Ratemap
Autocorr.
23.3Hz
11.7Hz
874 7072017 t14 c1
Lattice ratemap
Proj. ratemap
Proj. autocorr.
23.1Hz
Arena 1
Arena 2
Ratemap
Autocorr.
5.5Hz
2.9Hz
874 7072017 t14 c2
Lattice ratemap
Proj. ratemap
Proj. autocorr.
10.1Hz
Arena 1
Arena 2
Ratemap
Autocorr.
15.3Hz
15.5Hz
874 11072017 t13 c1
Lattice ratemap
Proj. ratemap
Proj. autocorr.
34.8Hz
Arena 1
Arena 2
Ratemap
Autocorr.
7.6Hz
9.9Hz
874 24072017 t13 c1
Lattice ratemap
Proj. ratemap
Proj. autocorr.
22.5Hz
Arena 1
Arena 2
Ratemap
Autocorr.
18.2Hz
18.9Hz
895 17112017 t16 c1
Lattice ratemap
Proj. ratemap
Proj. autocorr.
35.3Hz
Arena 1
Arena 2
Ratemap
Autocorr.
11.6Hz
6.7Hz
918 26032018 t9 c2
Lattice ratemap
Proj. ratemap
Proj. autocorr.
7.6Hz
Arena 1
Arena 2
Ratemap
Autocorr.
20.8Hz
21.3Hz
918 1042018 t9 c1
Lattice ratemap
Proj. ratemap
Proj. autocorr.
48.1Hz
Arena 1
Arena 2
Ratemap
Autocorr.
1.7Hz
1.8Hz
918 1042018 t9 c2
Lattice ratemap
Proj. ratemap
Proj. autocorr.
4.7Hz
Arena 1
Arena 2
Ratemap
Autocorr.
18.2Hz
17.1Hz
918 4042018 t9 c1
Lattice ratemap
Proj. ratemap
Proj. autocorr.
35.4Hz
Arena 1
Arena 2
Ratemap
Autocorr.
14.4Hz
9.6Hz
918 13042018 t9 c1
Lattice ratemap
Proj. ratemap
Proj. autocorr.
17.1Hz
Arena 1
Arena 2
Ratemap
Autocorr.
3.2Hz
2.8Hz
922 20022018 t16 c3
Lattice ratemap
Proj. ratemap
Proj. autocorr.
5.1Hz

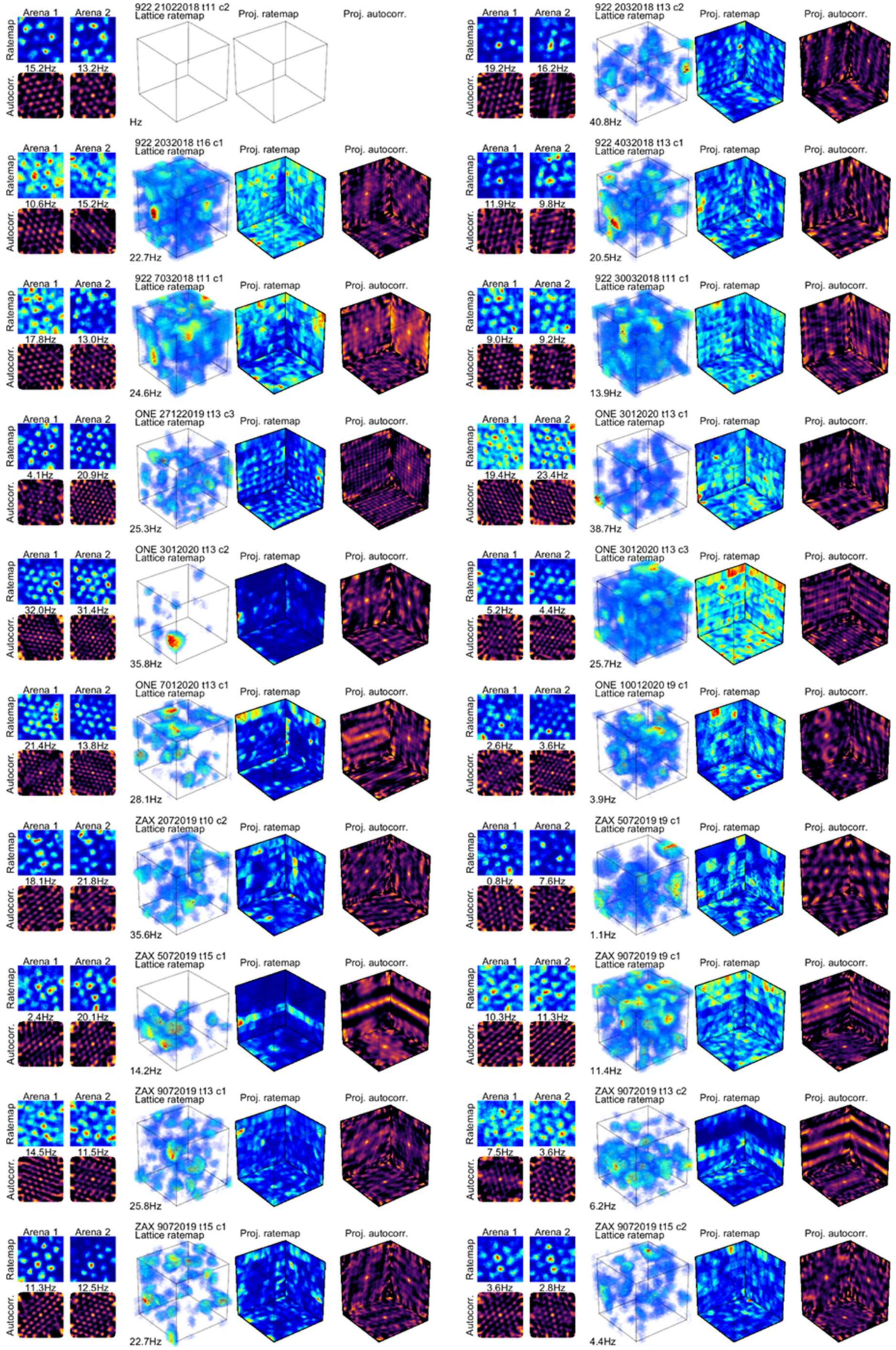
Arena 1
Arena 2
Ratemap
Autocorr.
Lattice ratemap
Proj. ratemap
Proj. autocorr.
922 21022018 t11 c2
15.2Hz
13.2Hz
Hz
922 2032018 t13 c2
19.2Hz
16.2Hz
40.8Hz
922 2032018 t16 c1
10.6Hz
15.2Hz
22.7Hz
922 4032018 t13 c1
11.9Hz
9.8Hz
20.5Hz
922 7032018 t11 c1
17.8Hz
13.0Hz
24.6Hz
922 30032018 t11 c1
9.0Hz
9.2Hz
13.9Hz
ONE 27122019 t13 c3
4.1Hz
20.9Hz
25.3Hz
ONE 3012020 t13 c1
19.4Hz
23.4Hz
38.7Hz
ONE 3012020 t13 c2
32.0Hz
31.4Hz
35.8Hz
ONE 3012020 t13 c3
5.2Hz
4.4Hz
25.7Hz
ONE 7012020 t13 c1
21.4Hz
13.8Hz
28.1Hz
ONE 10012020 t9 c1
2.6Hz
3.6Hz
3.9Hz
ZAX 2072019 t10 c2
18.1Hz
21.8Hz
35.6Hz
ZAX 5072019 t9 c1
0.8Hz
7.6Hz
1.1Hz
ZAX 5072019 t15 c1
2.4Hz
20.1Hz
14.2Hz
ZAX 9072019 t9 c1
10.3Hz
11.3Hz
11.4Hz
ZAX 9072019 t13 c1
14.5Hz
11.5Hz
25.8Hz
ZAX 9072019 t13 c2
7.5Hz
3.6Hz
6.2Hz
ZAX 9072019 t15 c1
11.3Hz
12.5Hz
22.7Hz
ZAX 9072019 t15 c2
3.6Hz
2.8Hz
4.4Hz

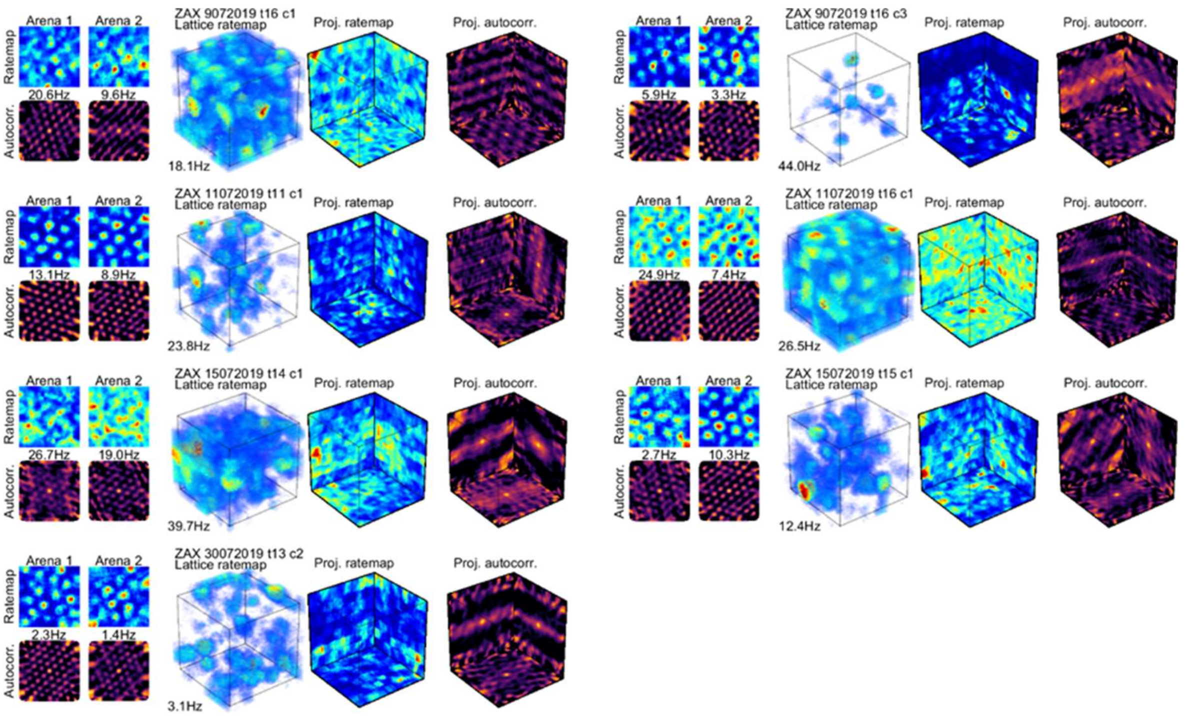


**Fig. S2. Arena and lattice firing rate maps for all analyzed grid cells.**

Colorbars can be seen in Fig. 1. Two grid cells are shown per row, with for each grid cell: Left) XY projected firing rate maps for the 1st and 2nd arena sessions respectively (top row) and autocorrelations of these (bottom row). Firing rate maps are scaled between 0Hz and the maximum firing rate value given beneath the map. Autocorrelations are scaled between -0.2 and 1. Middle left) the volumetric firing rate map in oblique view, voxels with a firing rate less than 10% of the maximum are transparent. Lines show the outermost edges of the lattice maze. Text above the map gives the rat number, date, tetrode and cluster of the cell. Text below the map gives the maximum firing rate displayed. Middle right) projected planar firing rate maps shown in their planes of projection. These are scaled between 0Hz and the maximum value in all 3 maps. Lines show the outermost edges of the lattice maze. Right) autocorrelations of the projected firing rate maps shown in their planes of projection. These are scaled between -0.2 and 1. Lines show the outermost edges of the lattice maze. It is unknown why cell 922 21022018 t11 c2 did not fire in the lattice maze but was recorded in both arena sessions.

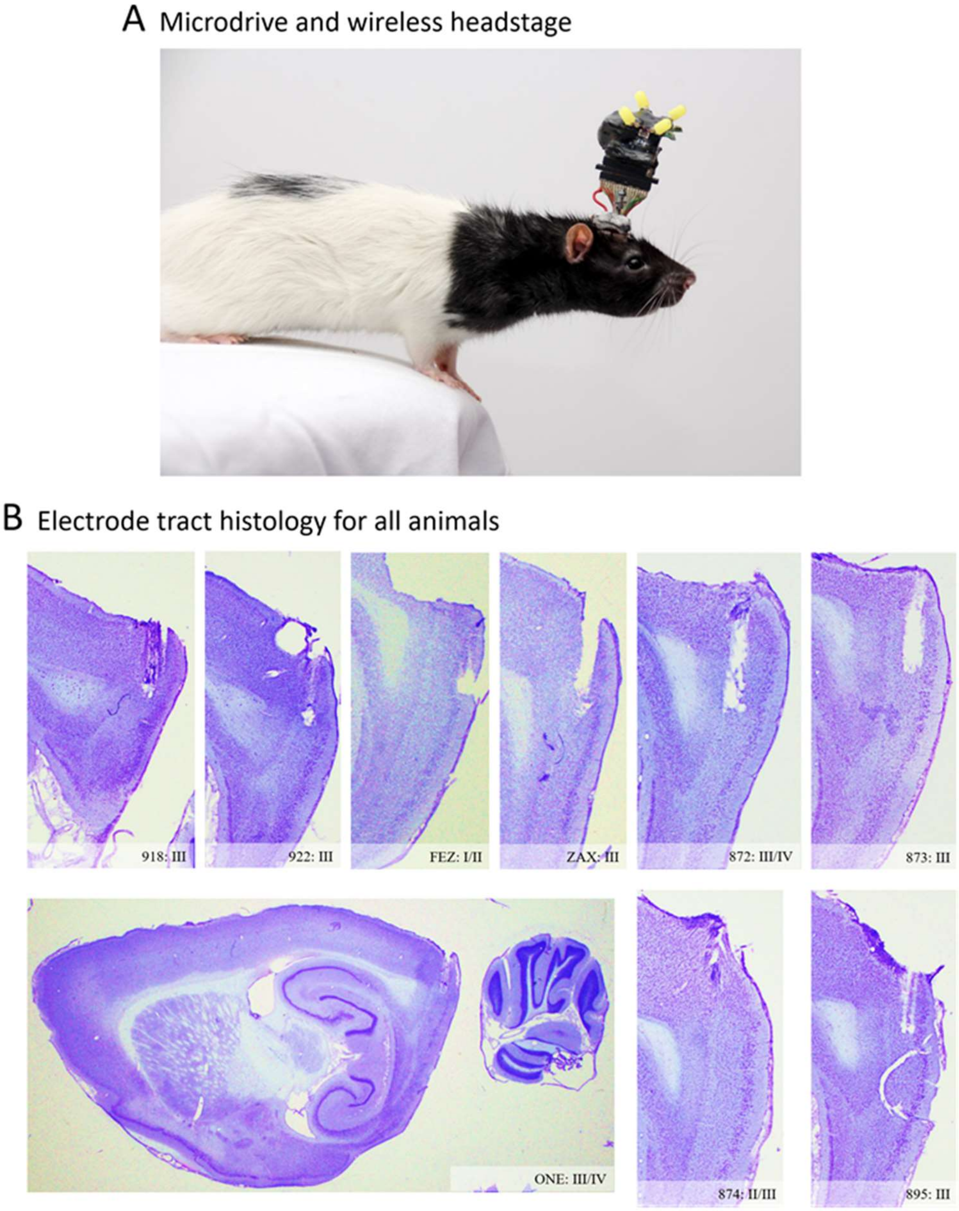


**Fig. S3. Electrodes mainly targeted layer III of the mEC.**

(A) Photograph of a rat implanted with a microdrive and connected to the wireless headstage. (B) Histological Nissl-stained sections. For each rat, the sagittal brain section best demonstrating the position of the electrodes is given, text gives the animal identification and estimated layer of the mEC where recordings were made. Close-ups of the mEC are provided for every rat except rat ‘ONE’ where the whole brain section is given.

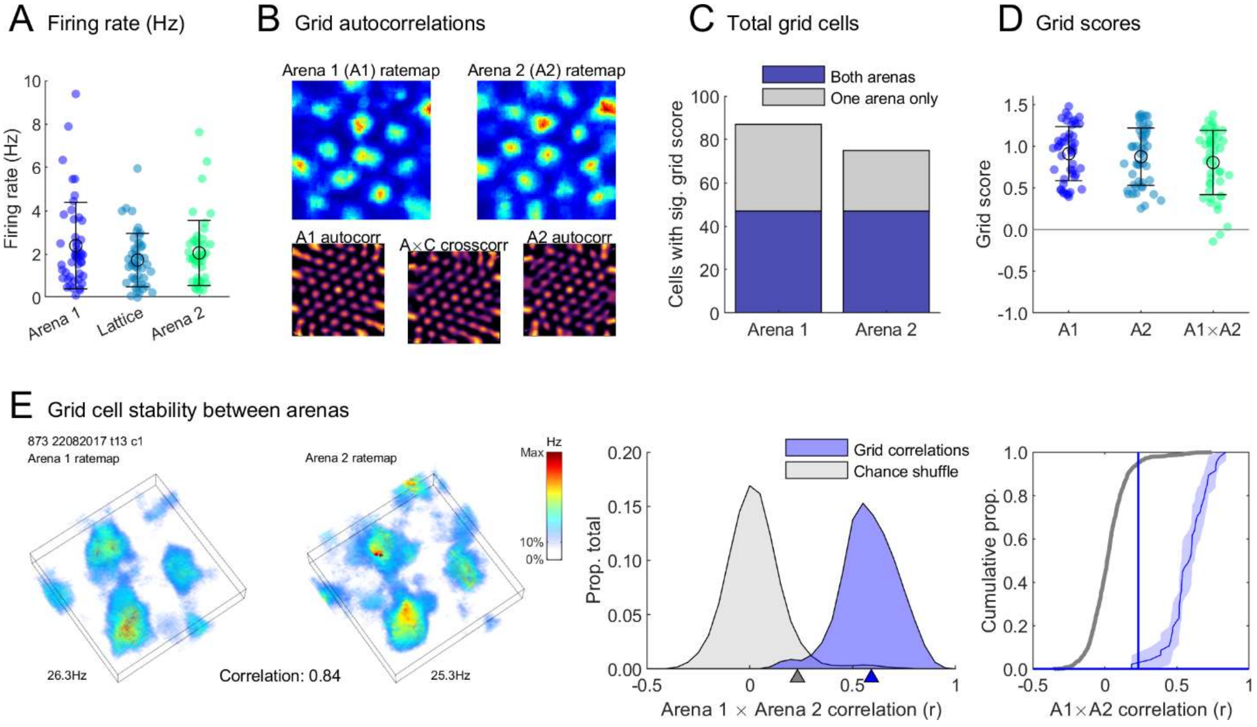


**Fig. S4. Grid cells were more stable than chance throughout recordings.**

(A) Grid cell firing rates did not differ between the mazes ($F(2,138) = 2.0$, $p = .135$, $\eta^2 = 0.029$, one-way ANOVA). (B) Examples autocorrelations in the arena sessions. In addition to ratemap autocorrelations we also calculated the cross-correlation between arenas 1 and 2. Only cells with a grid score exceeding the 95$^{th}$ percentile of a shuffle in both arena sessions were analysed. (C) Total number of cells with a grid score exceeding the 95$^{th}$ percentile of a shuffle in each arena session. The purple bar sections represent the cells categorised as grid cells and included in the main analyses. (D) Grid scores of all grid cells calculated for both arenas and the cross-correlation between arena maps. These scores did not differ ($F(2,138) = 1.1$, $p = .337$, $\eta^2 = 0.016$, one-way ANOVA). (E) Left) example firing of a grid cell in both arenas. Middle) to determine if grid cells were stable between arenas we correlated their arena firing rate maps (blue area; blue triangle denotes median) and compared this distribution to correlations between 5000 shuffled ratemaps (grey area; grey triangle denotes 95$^{th}$ percentile). Right) cumulative density curves of the same distributions. The blue vertical line marks the 95$^{th}$ percentile of the shuffle on the x-axis and the blue horizontal line marks the y-intercept of the observed correlation distribution with this. Grid cells were more stable than chance (shuffled arena correlations 95$^{th}$ percentile: 0.23, grid cell median correlation: 0.59, D = 0.96, $p < .001$, two-sample Kolmogorov-Smirnov test).

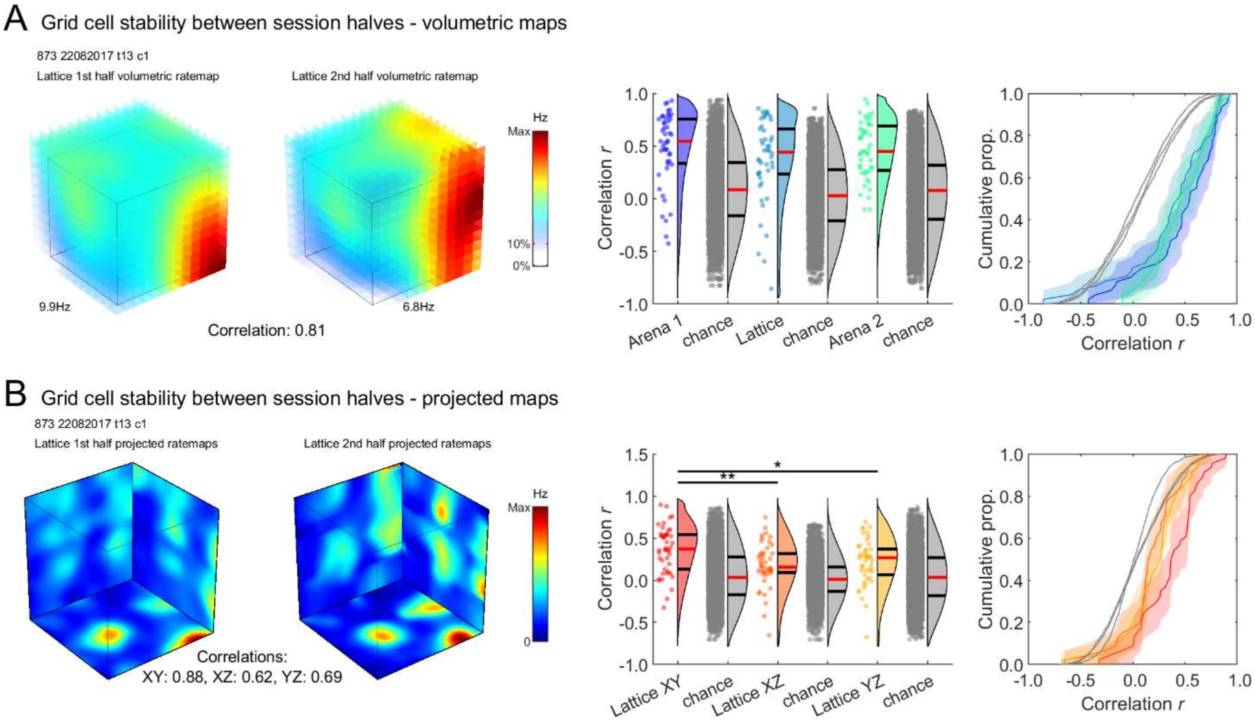


**Fig. S5. Grid cells were more stable than chance within sessions, especially in the XY plane.** See Methods: *Spatial stability within sessions*. (A) Left) representative grid cell volumetric firing rate maps used for the volumetric correlation analysis; each represents one half of the same lattice session. Middle) raincloud plots showing the distribution of correlation values found for the arena and lattice sessions. Grey distributions represent correlations between random cells recorded in each maze. Red lines denote medians, black lines denote 1$^{st}$ and 3$^{rd}$ quantiles. Right) cumulative distribution functions of the same distributions. In all sessions grid cells were more stable than chance ($p < .0001$ in all cases; two-sample t-tests) and the mazes did not differ ($F(2,137) = 1.6$, $p = .203$, $\eta^2 = 0.023$; one-way ANOVA). (B) Left) same cell as A but shown using projected maps used for planar correlations. Middle) raincloud plots as in A, showing the correlations found for each projected plane of the lattice. Right) cumulative distribution functions of the same distributions in A. All projections were more stable than chance ($p < .001$ in all cases; two-sample t-tests) but horizontal (XY) projections yielded significantly higher correlations than vertical ones ($F(2,135) = 5.8$, $p = .004$, $\eta^2 = 0.079$, lattice XY vs XZ or YZ $p < .04$, all other $p > .05$; one-way ANOVA).

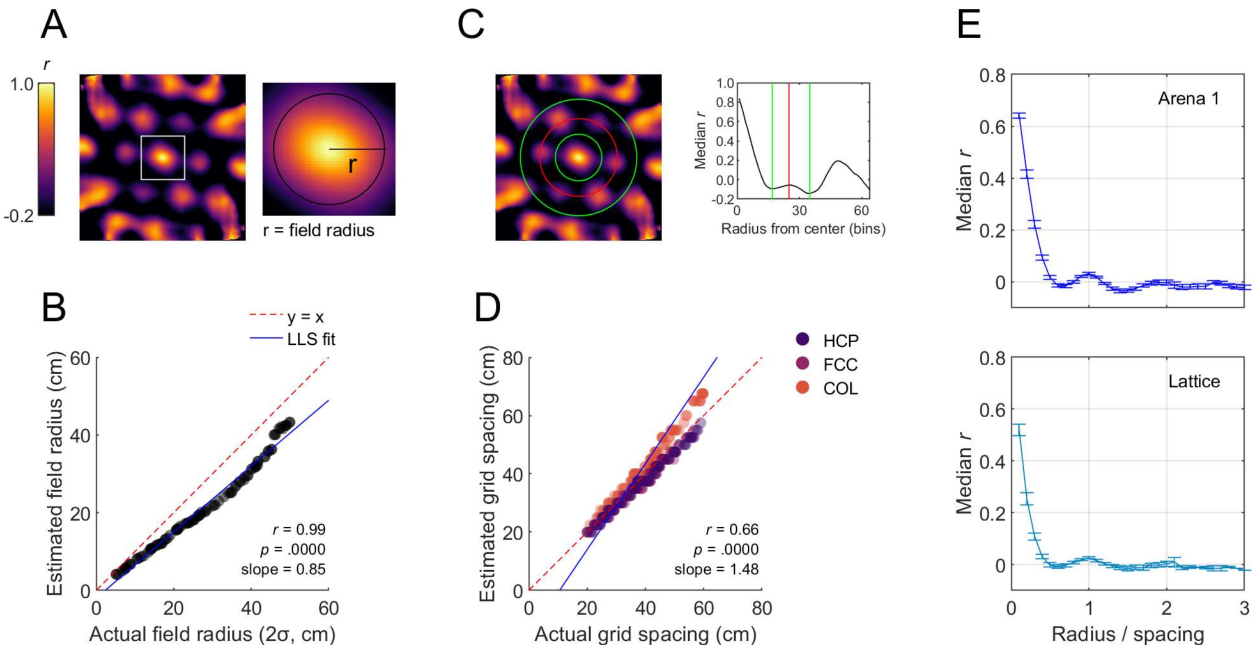


**Fig. S6. Schematics and verification of the field radius and grid spacing analyses**

(A) Two-dimensional schematic showing the field size (radius) estimation procedure. The autocorrelation (top left) was thresholded at 0.25 and field size was estimated as the radius of the remaining central peak. (B) This analysis was validated on simulated FCC field arrangements with varying field sizes (100 arrangements with field sigmas between 1 and 10 bins and grid spacing in mm equal to 120σ; see Methods: *Simulated field arrangements*). Note that if fields overlap this analysis will tend to overestimate field size. (C) Two-dimensional schematic showing the field spacing (scale or wavelength) estimation procedure. The median correlation found at different distances from the central peak were mapped and grid spacing was estimated as the location of the first peak after the central one. (D) This analysis was validated on HCP, FCC and columnar (COL) field arrangements (random fields do not have an 'actual' spacing to validate against). Note that the size of our mazes limited field spacing to a maximum of approximately 120cm. (E) The field spacing estimation results from real grid cells. For each cell the spacing was estimated as in C, shown here is the median correlation at every radius normalized by the first peak (the estimated spacing). A clear peak and periodicity can be seen in both the arena (top) and lattice (bottom). Cells for which no spacing could be estimated (0% in the arena, 40.2% in the lattice) are not shown.

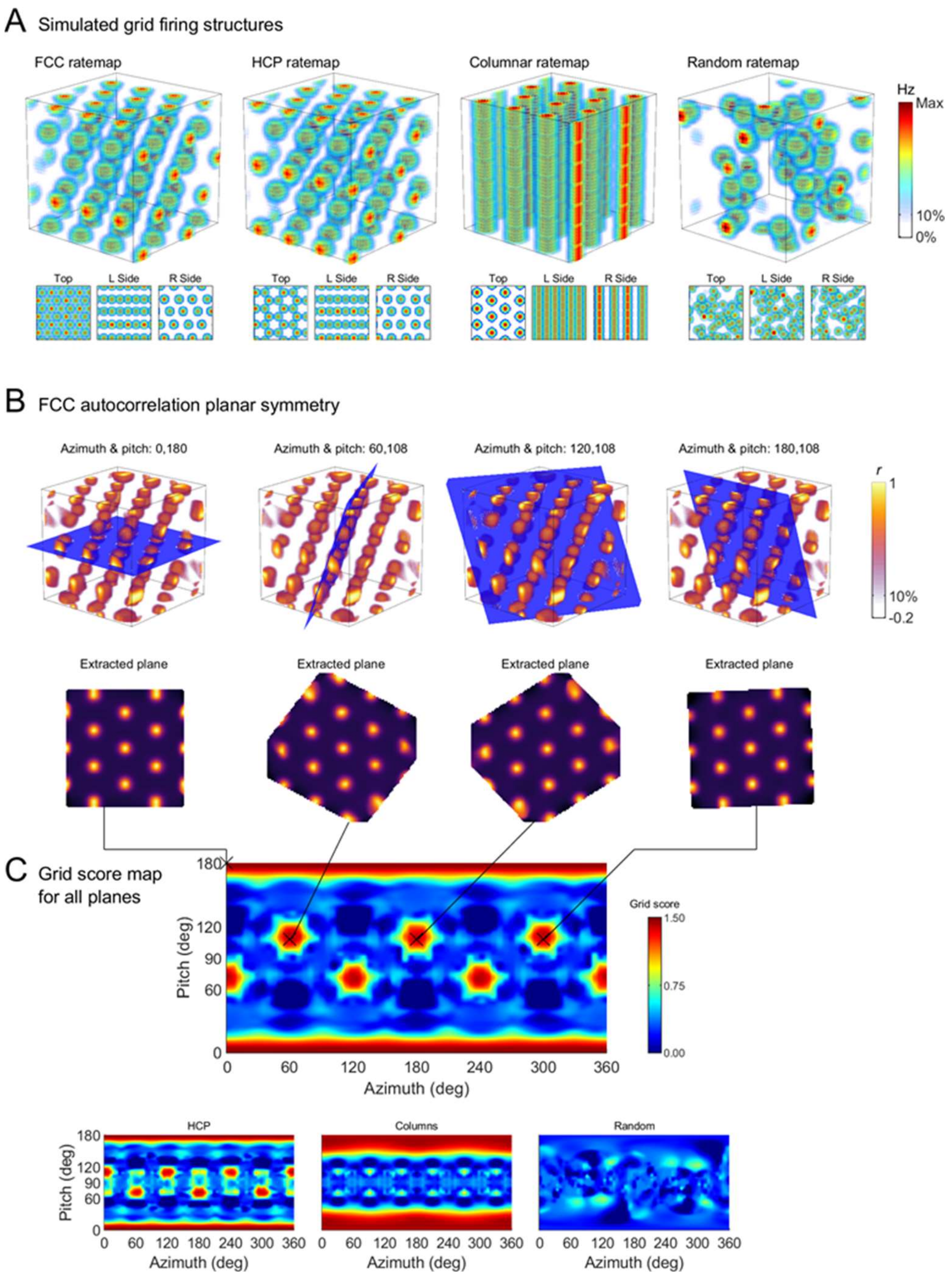


**Fig. S7. Schematic demonstrating the planar symmetry analysis.**

(A) Example simulated field arrangements in oblique and side views. (B) Taking the autocorrelation of the FCC arrangement shown in A, it is clear there are four planes that can be cut through the centre (top row) which slice through fields arranged in a planar hexagonal pattern (bottom row). (C) We extract all possible planes and calculate the hexagonal grid score for each one. Mapping these according to their pitch/azimuth yields different patterns for the different field arrangements which can be used to differentiate them.

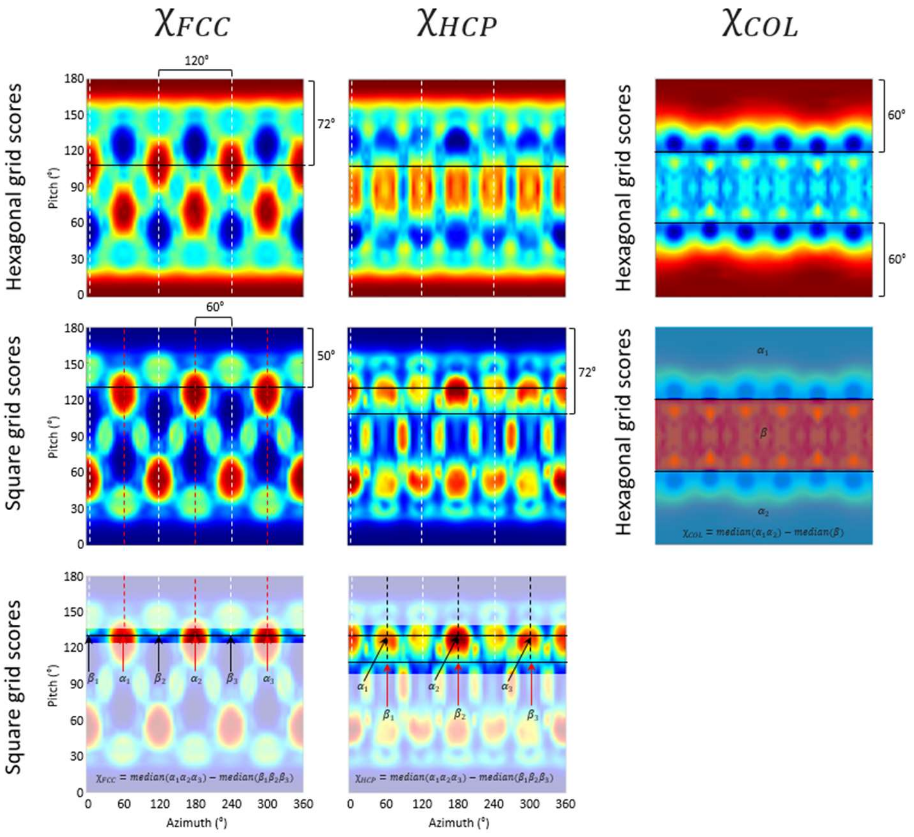


**Fig. S8. Schematic showing how the structure scores ($\chi_{FCC}$, $\chi_{HCP}$ & $\chi_{COL}$) were calculated.** After extracting all possible planes through a cell's spatial autocorrelation we calculated the hexagonal and square grid score for each one. Mapping these according to their pitch/azimuth yields different patterns for the different field arrangements which can be used to differentiate them. In each example the grid score maps have already been corrected so that the 'best plane' is horizontal (placing the highest hexagonal grid scores at the top and bottom of the maps). Left column demonstrates the process for the $\chi_{FCC}$ score: first, the three values at 72° from the best plane and separated by 120° between them with the maximum total sum were found. In an FCC arrangement, low square grid scores are found at 50° to the best plane at the same azimuthal angles and high square grid scores can be found offset 60° from them in azimuth. Thus, $\chi_{FCC}$ was calculated as the difference between these points. Middle column demonstrates the process for the $\chi_{HCP}$ score: again, the three values at 72° from the best plane and separated by 120° between them with the maximum total sum were found. In an HCP arrangement, high square grid scores are found at 50° to the best plane and offset 60° from them in azimuth, while low square grid scores are found at 72° to the best plane and at the same azimuth angles. Thus, $\chi_{HCP}$ was calculated as the difference between these points. Right column demonstrates the process for the $\chi_{COL}$ score: in a columnar arrangement there is a large field of pitch angles that intersect the columns and exhibit a high hexagonal grid score, however, there are virtually no high grid scores perpendicular to this plane. Thus, $\chi_{COL}$ was calculated as the difference between the median grid score found at the angles 0 - 60° & 120 - 180° and the median score found at the angles 60 to 120°. These angles were chosen as they divide the spherical domain into two regions with equal surface area (the poles and an equatorial 'belt').

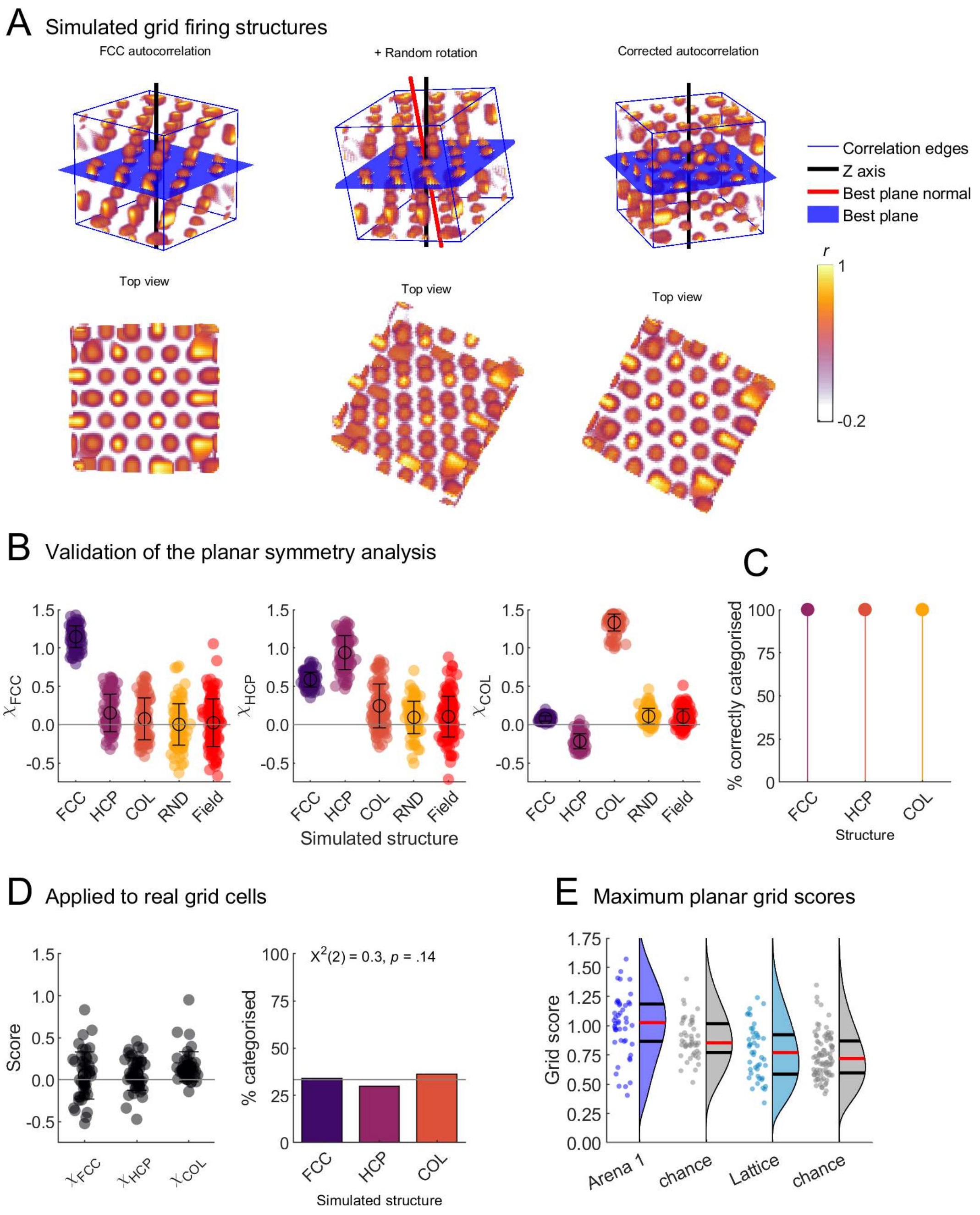


**Fig. S9. Explanation and validation of the planar symmetry analysis.**

(A) Autocorrelations were randomly rotated before analysis by the planar symmetry method to avoid assumptions of the structure's orientation. This schematic shows the correct identification of the 'best plane' or the plane with the highest grid score and successful reorientation of the autocorrelation based on this. (B) Left) for every simulated field arrangement we calculated the three symmetry scores ($\chi_{FCC}$, $\chi_{HCP}$ and $\chi_{COL}$), in each case the correct score is highest for its corresponding field arrangement while the random ('RND') and shuffled field ('Field') arrangements are low in all three scores. (C) If simulated cells are categorized based on these scores (category = configuration with max score) they are grouped with 100% accuracy, confirming that the planar symmetry analysis can correctly identify field arrangements. (D) Left) the three symmetry scores found for the real grid cells, all scores are much lower than expected if the arrangements were present and there was no significant difference between them ($F(2,138) = 2.9$, $p = 0.0584$, $\eta^2 = 0.040$, one-way ANOVA). Right) if grid cells are categorized as in C an equivalent proportion of cells fall into each group meaning that no one field arrangement dominates the data. (E) For each grid cell we calculated the maximum possible planar grid score (the maximum value found in the hexagonal grid score maps in Fig. S8) and compared these to shuffled arrangements of firing fields (2 per cell; Methods: *Grid field shuffle*). Arena grid scores were higher than chance ($t(138) = 2.7$, $p = .0072$, Cohen's $d = 0.47$), while lattice scores were not different from chance ($t(135) = 0.9$, $p = .3668$, Cohen's $d = 0.16$) suggesting that there were not hexagonal arrangements of fields on any plane through the lattice.

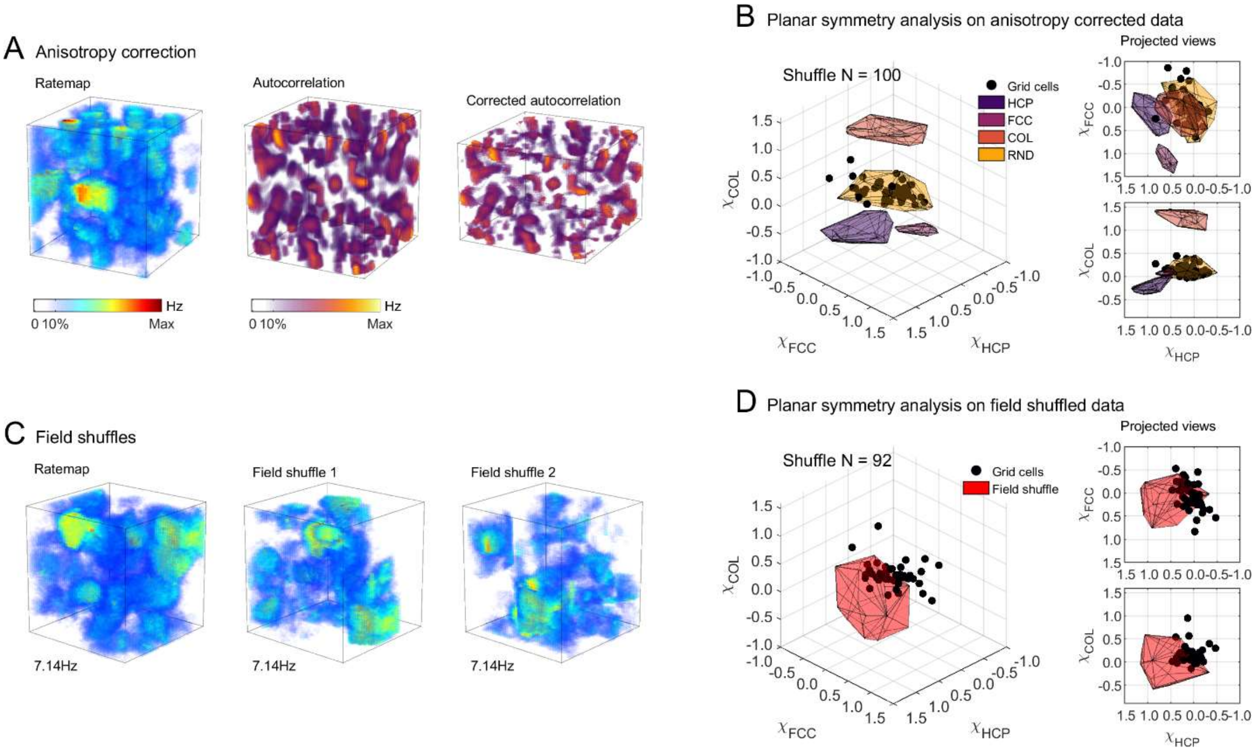


**Fig. S10. Planar symmetry analysis yields similar results using a field shuffle.**

(A) We repeated the planar symmetry analysis after correcting grid cell autocorrelations for anisotropy (Methods: *Autocorrelation correction for anisotropy*). An example of a grid cell ratemap is shown (left), the autocorrelation of this ratemap (middle) and the same autocorrelation after correcting for field anisotropy (right). Note that the corrected autocorrelation is compressed along the vertical axis and that the central peak is more spherical than in the uncorrected version. (B) The configuration scores for these autocorrelations did not differ greatly from uncorrected ones. (C) In addition to the FCC, HCP, columnar and random field arrangements described elsewhere we also replicated the grid field shuffle described by Barry and Burgess (*25*) which randomly shuffles field positions while maintaining their local firing activity as best as possible. Owing to the computational demands of this analysis we performed the shuffle only twice per grid cell (92 shuffles). An example of a grid cell ratemap is shown on the left and the same ratemap with shuffled firing fields on the right. (D) The ratemaps in this shuffled dataset exhibit some higher FCC and HCP scores (red polygon) than the real grid cells (black markers). However, on average the scores are all centred on zero, like the real grid cells (values shown in Fig. S9B, 'Field' group).

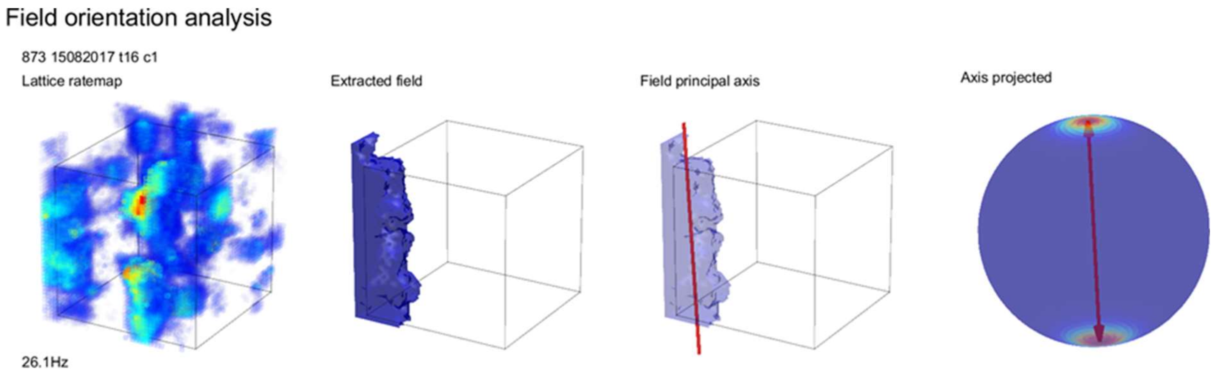


**Fig. S11. Schematic demonstrating the field orientation analysis.**

An example firing rate map (left) is thresholded at 20% of the peak firing rate and regions which passed our criteria were considered place fields (2nd plot shows an example field). We can visualize these regions as convex hulls (3rd plot) and extract features such as their principal axis (red line). To visualize the orientation of multiple fields we project these axes onto a unit sphere and generate a spherical Von-Mises kernel smoothed density map, where hot colors denote that many fields 'pointed' their principal axis in this orientation.

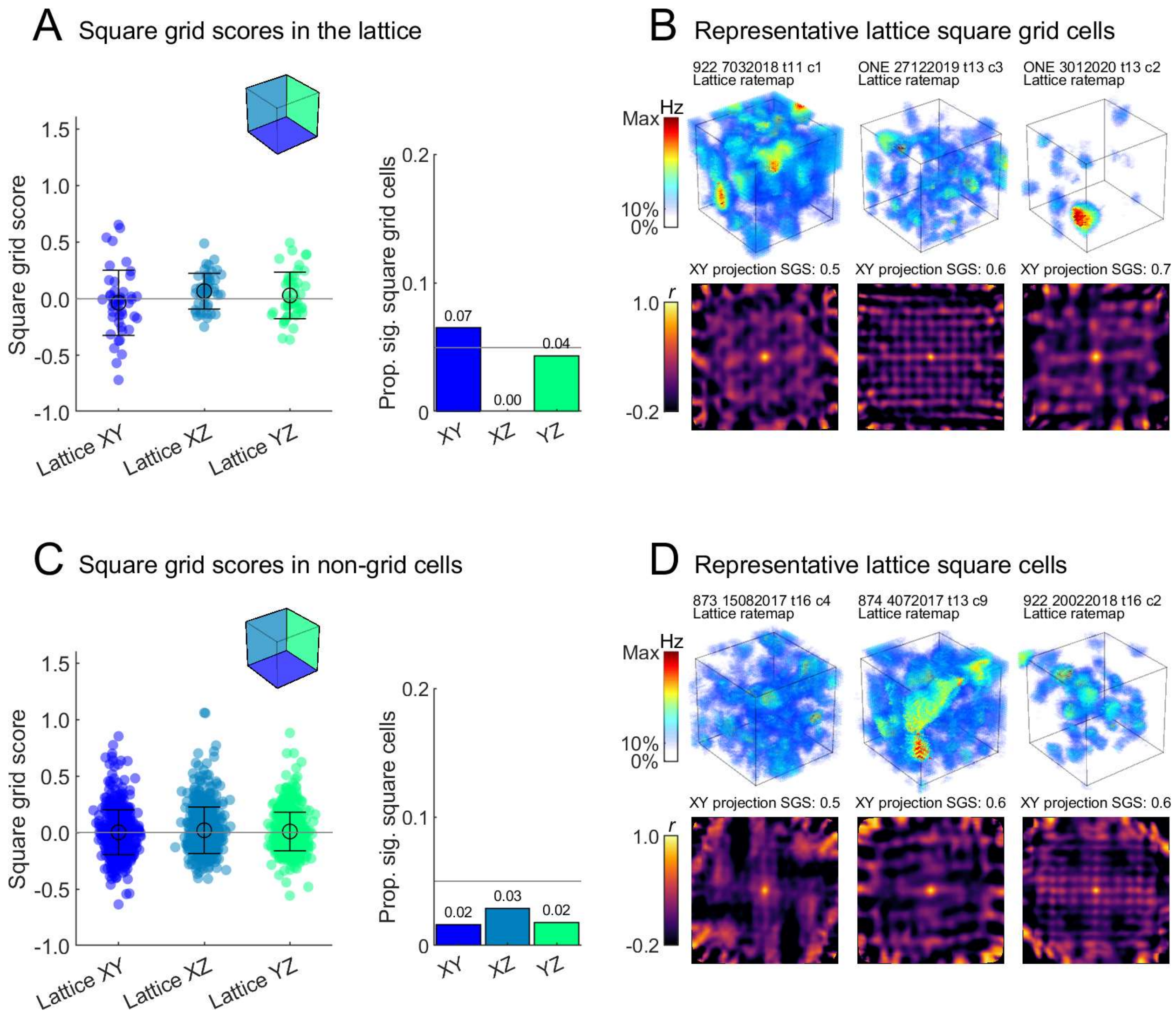


**Fig. S12. Weak evidence of square pattern activity in grid cells and non-grid cells.**

(A) Left) square grid scores for all grid cells in each projected plane of the lattice maze. Square grid scores are low in each of the lattice planes and these did not differ ($F(2,135) = 2.5$, $p = .0872$, $\eta^2 = 0.036$, one-way ANOVA). Right) proportion of grid cells with a square grid score exceeding the 95th percentile of a chance distribution in each lattice plane. A small number of cells exhibited a significant square firing pattern when projected onto the XY and YZ planes. Grey line shows the value that would be expected by chance (5%). (B) Three examples of these XY square grid cells. Top row shows the volumetric firing rate map, text gives the rat number, date, tetrode and cluster. Bottom row shows the autocorrelation of the XY projected firing rate map, text gives the square grid score (SGS) of the autocorrelation. (C-D) same as A-B but for all non-grid cells. Square grid scores were again low and did not differ between projected planes ($F(2,1648) = 1.2$, $p = .2975$, $\eta^2 = 0.0015$, one-way ANOVA). No plane exhibited more square grid cells than would be expected by chance (5%, grey line).

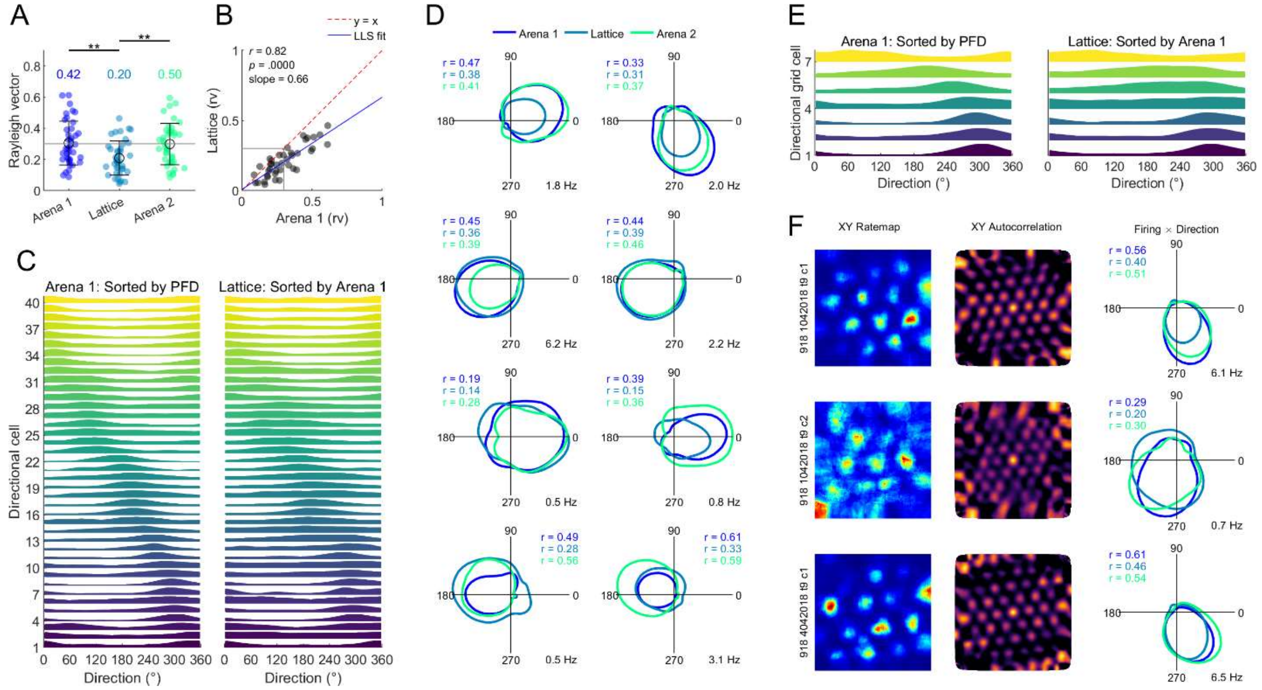

**Fig. S13. Directional modulation was preserved in the lattice.**

These analyses were conducted on head direction estimated from speed filtered displacement. (A) Rayleigh vector lengths for all 40 directionally modulated cells (i.e., any cells with a Rayleigh vector length exceeding the 95th percentile of 100 spike train shuffles in both arena sessions, may include grid cells). Grey line denotes a commonly used arbitrary cut-off (0.3) and text gives the proportion of cells with a Rayleigh vector length greater than this. Vector lengths were lower in the lattice ($F(2,117) = 7.1$, $p = .0013$, $\eta^2 = 0.108$, lattice vs arena 1 or 2 $p < .004$, all other $p > .05$, one-way ANOVA). (B) Vector lengths were correlated between arena 1 and the lattice suggesting preserved directional modulation in this maze. (C) Left) normalized arena 1 tuning curves for all directionally modulated cells sorted by their preferred firing direction (PFD; peak firing). Right) normalized lattice tuning curves sorted by their arena 1 PFD. The preserved diagonal ordering suggests that cells maintained the same allocentric firing directions in the lattice; the population correlation between arena and lattice tuning curves ($r = 0.85$) exceeded the 95th percentile of 1000 shuffles ($r = 0.33$, z = 8.3) confirming this. (D) Overlaid arena and lattice polar tuning curves for 8 example directional cells. Coloured text gives the Rayleigh vector length for the corresponding tuning curves. (E) Same as C but for all conjunctive grid × direction cells (n = 7). As before, the correlation between arena and lattice tuning curves ($r = 0.91$) exceeded the 95th percentile of 1000 shuffles ($r = 0.55$, z = 3.7) confirming that conjunctive grid cells maintained the same allocentric firing directions in the lattice. (F) Example conjunctive grid cells, one per row, ratemap, spatial autocorrelation and overlaid tuning curves as in D.

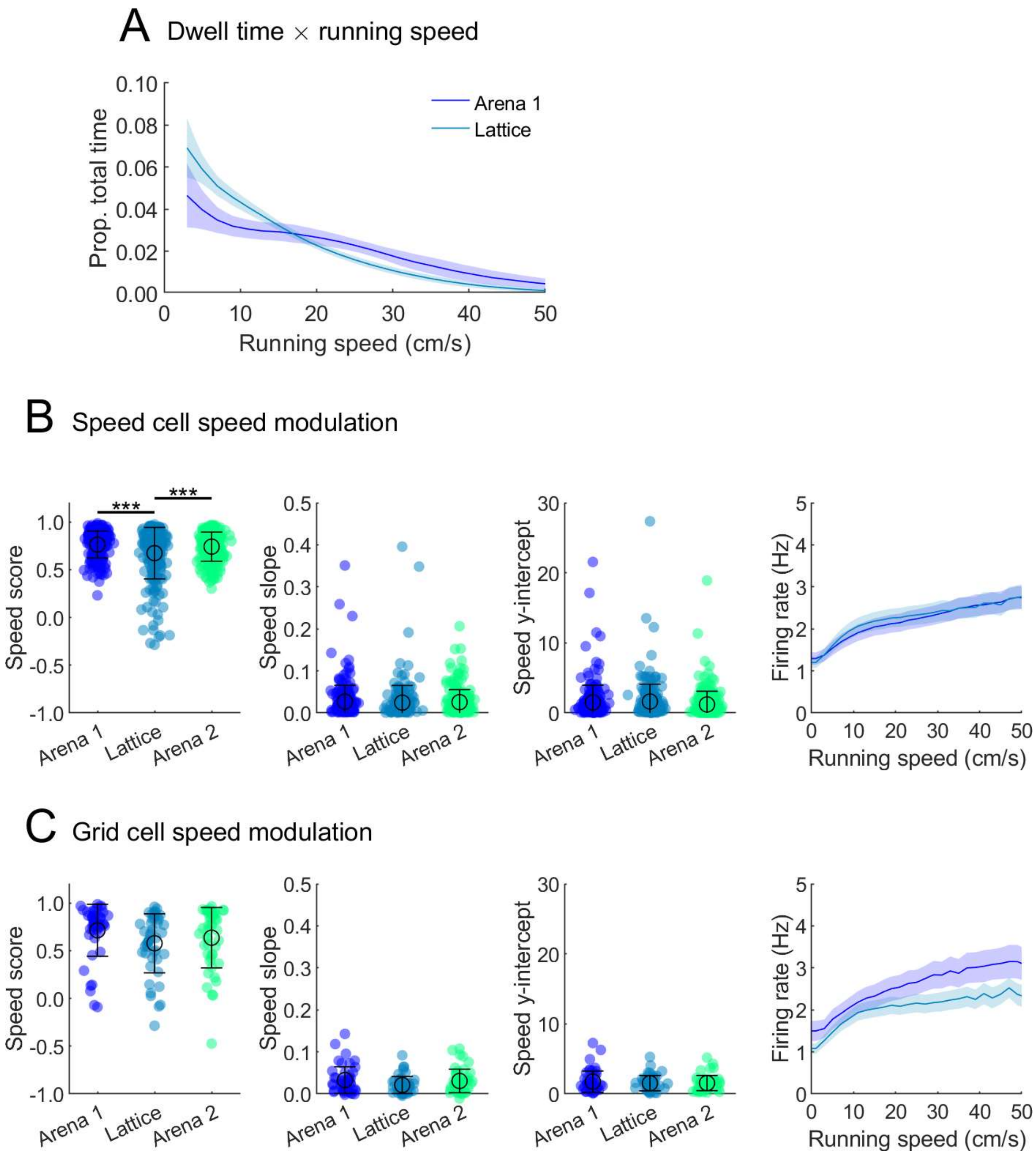


**Fig. S14. Very little change in speed cell activity and speed coding by grid cells.**

(A) The mean and SEM proportion of time animals spent moving at various running speeds, averaged across sessions. (B) Speed scores of speed modulated non-grid cells dropped in the lattice ($F(2,663) = 12.6$, $p < .0001$, $\eta^2 = 0.0367$, one-way ANOVA), however, the slope and y-intercept of speed-firing rate relationships were unaffected (all $p > .10$, one-way ANOVAs). (C) Speed scores, slope and y-intercepts of speed-firing rate relationships were also unaffected in grid cells (all $p > .08$, one-way ANOVAs).

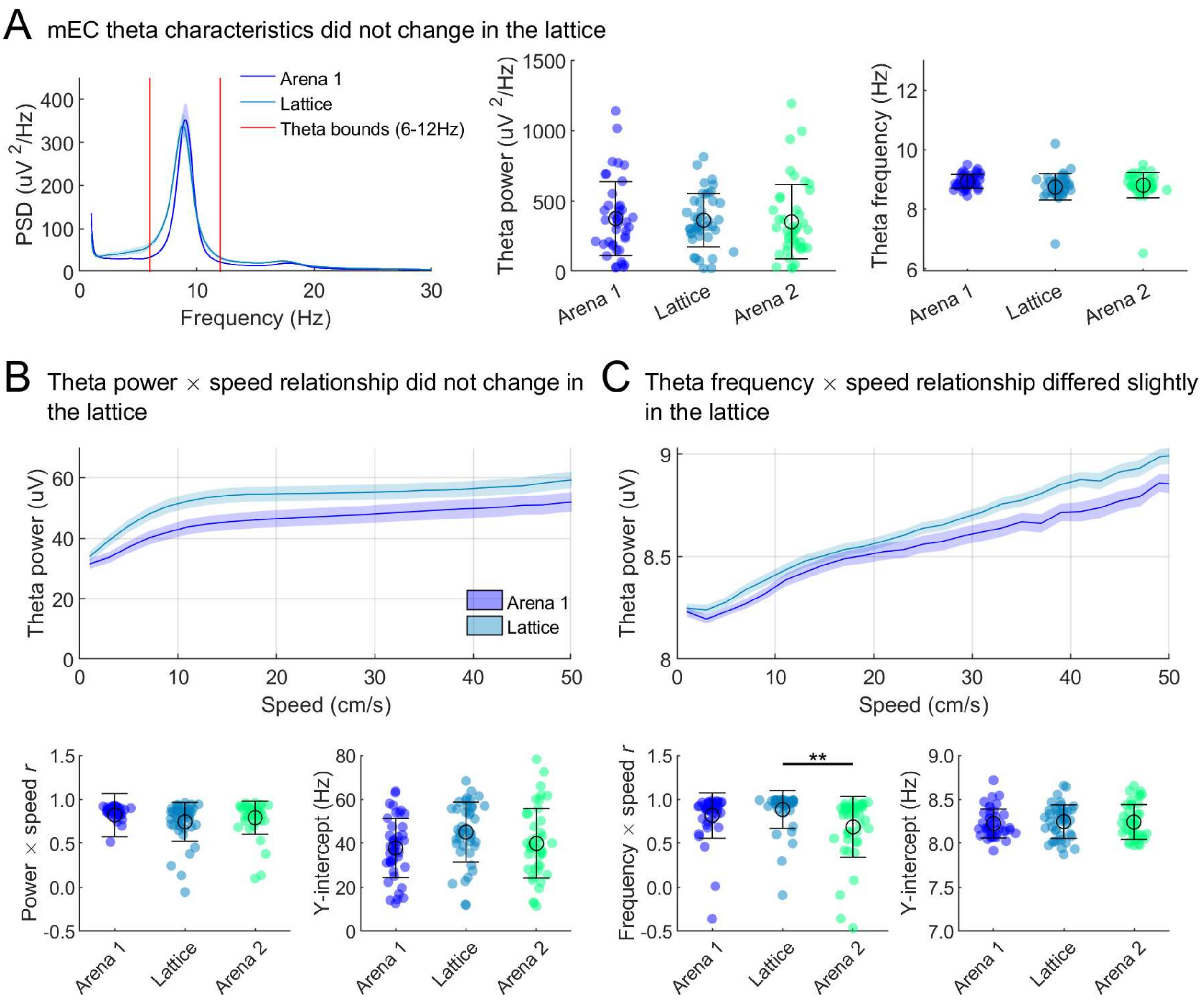


**Fig. S15. Very little change in theta power and frequency or relationship with running speed.**

Markers represent sessions, averages and errors computed across sessions. (A) Left) the mean PSD in each maze, averaged across sessions. Red lines denote the frequency band associated with theta rhythm. Middle) maximum power associated with the theta band in each session, these did not differ ($F(2,123) = 0.1$, $p = .9180$, $\eta^2 = 0.0014$, one-way ANOVA). Right) frequency associated with max theta power in each maze session, these did not differ ($F(2,123) = 2.5$, $p = .0828$, $\eta^2 = 0.0397$). (B) Mean and SEM theta power (amplitude of the Hilbert transform) observed at various running speeds, averaged across sessions. Below this are boxplots where markers represent sessions. These show the Pearson's correlation between power and speed which did not differ between mazes (left; $F(2,123) = 1.2$, $p = .2957$, $\eta^2 = 0.0196$, one-way ANOVA) and the y-intercept of the power and speed relationship which also did not differ (right; $F(2,123) = 2.9$, $p = .0602$, $\eta^2 = 0.0447$, one-way ANOVA). (C) Same as B but for theta frequency. The correlation between theta frequency and speed decreased in the second arena session with respect to the lattice ($F(2,123) = 5.6$, $p = 0.0046$, $\eta^2 = 0.0839$, lattice vs arena 2 $p = .0037$, all other $p > .05$, one-way ANOVA) while the y-intercept did not differ (right; $F(2,123) = 2.9$, $p = .8367$, $\eta^2 = 0.0029$, one-way ANOVA).

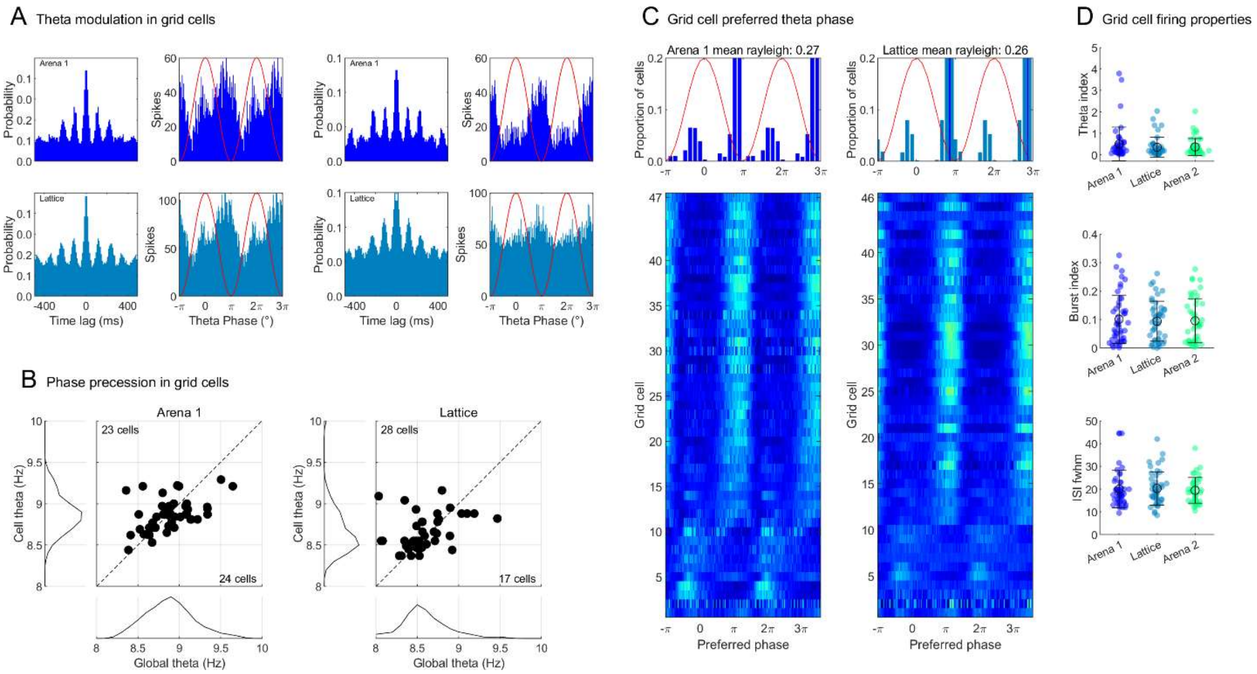


**Fig. S16. Grid cell spiking dynamics were preserved in the lattice maze.**

(A) Spike probability properties of two example grid cells. Left four plots are for one cell, right four plots are a different cell. Top row gives data for the first arena, bottom row gives data for the lattice. For each maze the left plot shows the 400ms spike autocorrelogram (1ms bins) and the right plot shows the spike-theta phase histogram. Red lines trace the amplitude of a scaled theta wave at every phase. Theta modulation and phase preference were unaffected in the lattice. (B) Scatter plots showing the frequency of intrinsic theta (extracted from the spike autocorrelograms) vs extrinsic theta (extracted from the power spectral density estimate of the LFP). Markers represent grid cells, numbers represent counts that are above or below the diagonal. Markers above the dashed line represent cells firing at a rate faster than global theta, an indicator of phase precession. Cells were not modulated at a higher frequency than extrinsic theta in the arena ($\chi^2(1) = 0.02$, $p = .8840$) or lattice ($\chi^2(1) = 2.69$, $p = .1011$, Chi-square tests of expected proportions). (C) Histograms showing the preferred theta phase (value of circular mean) of all grid cells. Red lines trace the amplitude of a scaled theta wave at every phase, blue lines show the circular mean of the preferred phases. Cells showed an equal preference for one theta phase in both mazes ($F(1,92) = 0.1$, $p = .7067$, $\eta^2 = 0.0016$, one-way ANOVA comparing spike phase Rayleigh vector lengths) and these preferred phases did not differ between the mazes (k = 423, $p > .99$, two-sample Kuiper test). Shown below each plot is the firing probability of all grid cells relative to theta phase. Each row represents a cell and rows are sorted from top to bottom by the circular mean of the cell's phase angles from early to late. (D) General firing properties such as the theta index extracted from each grid cell's spike autocorrelation (top; $F(2,135) = 0.9$, $p = .3905$, $\eta^2 = 0.0138$, one-way ANOVA), burst index (middle; $F(2,137) = 0.1$, $p = 0.9025$, $\eta^2 = 0.0015$, one-way ANOVA) and the full width at half maximum of the interspike interval (fwhm of the ISI: bottom; $F(2,130) = 0.1$, $p = 0.8616$, $\eta^2 = 0.0023$, one-way ANOVA) were unchanged in the lattice maze.

**Table S1. Sessions cell statistics for each rat**

| | Rat | Sessions | Cells | Grid cells (% cells) | Directional cells (% cells) | Speed cells (% cells) | mEC layer |
|---|---|---|---|---|---|---|---|
| | 872* | 2 | 65 | 0 (0.0) | 0 (0.0) | 9 (4.0) | 3&4 |
| | 873[a] | 3 | 82 | 8 (9.8) | 0 (0.0) | 25 (11.3) | 4 |
| | 874 | 4 | 64 | 5 (7.8) | 3 (7.5) | 20 (9.0) | 2&3 |
| | 895 | 2 | 48 | 1 (2.1) | 1 (2.5) | 21 (9.5) | 3 |
| | 918 | 5 | 26 | 5 (19.2) | 19 (47.5) | 14 (6.3) | 3 |
| | 922 | 10 | 97 | 7 (7.2) | 10 (25.0) | 47 (21.2) | 3 |
| | FEZ* | 3 | 68 | 0 (0.0) | 0 (0.0) | 7 (3.2) | 1&2 |
| | ONE | 5 | 70 | 6 (8.6) | 0 (0.0) | 32 (14.4) | 3&4 |
| | ZAX | 8 | 105 | 15 (14.3) | 7 (17.5) | 47 (21.2) | 3 |
| Total | 9 | 42 | 625 | 47 (7.5) | 40 (6.4) | 222 (35.5) | - |

Session and cell totals for each rat. Histological verification of the mEC layers can be seen in Fig. S3.

*Note that two rats did not yield any grid cells but contributed to the behavior and other electrophysiological data.

[a]Rat 873 exhibited the horizontal planar grid firing patterns.

**Movie S1.**

Representative grid cell activity in the lattice maze from different animals and sessions - 3D rotating spike and position plots with volumetric ratemaps, projected ratemaps and autocorrelations. Two-dimensional arena firing rate maps are also shown.
